# A survey of pairwise epistasis supports an outside-in hierarchy of clade-specifying and function-defining residues in PSD95 PDZ3

**DOI:** 10.1101/2020.06.26.174375

**Authors:** David Nedrud, Willow Coyote-Maestas, Daniel Schmidt

## Abstract

Deep mutational scanning enables data-driven models of protein structure and function. Here, we adapted Saturated Programmable Insertion Engineering as an economical and programmable deep mutational scanning technique. We validate this approach with an existing single mutant dataset in the PSD95 PDZ3 domain, and further characterize most pairwise double mutants to study how a mutation’s phenotype depends on mutations at other sites, a phenomenon called epistasis. We observe wide-spread proximal negative epistasis, which we attribute to mutations affecting thermodynamic stability, and strong long-range positive epistasis, which is enriched in an evolutionarily conserved and function-defining network of ‘sector’ and clade-specifying residues. Conditional neutrality of mutations in clade-specifying residues compensates for deleterious mutations in sector positions. This suggests an outside-in hierarchy of interactions through which positive epistasis between clade-specifying residues and the PDZ sector facilitated the evolutionary expansion and specialization of PDZ domains.

## Introduction

A protein’s primary sequence encodes its structure, conformational dynamics, and function. Mutations to this sequence are informative perturbations because they provide access to emergent protein properties that arise from the collective physical interactions of all amino acids within a protein. These perturbations, particularly from higher-order mutations, are difficult to predict. Thus, experimentally measuring perturbations from mutations provide crucial insight into biochemical mechanisms of protein function such as enzyme catalysis and ligand binding. Mutations allow us to map how residues that contribute to these functions are distributed in a protein’s tertiary and quaternary structure, and to identify determinants of protein folding and stability. High-throughput mutagenesis techniques, phenotyping assays, and sequencing enable deep mutational scanning (DMS) ^1^ in which the impact of replacing every residue of a protein with all 19 alternative amino acids is measured. DMS thus facilitates data-driven models of protein structure and function, which provide insight into enzyme activity, protein binding fitness landscapes ^2–9^, improve rational protein engineering ^10^, and functional genomics-guided oncology^11–13^.

Naturally occurring mutations result in variation, which is the raw material of evolutionary processes. In experimental evolution, mutations are useful to probe the molecular and mechanistic basis of adaptation. Interactions between multiple mutations shape and constrain evolutionary pathways of proteins; this dependence of a mutation’s phenotype on mutations at other sites is called epistasis ^14–16^. Epistasis plays a key role in protein evolvability and robustness by increasing the number of viable mutational trajectories that sidestep deleterious intermediates ^16^. In pioneering work, DMS was applied to map global epistasis on the IgG-binding domain of protein G (GB1) ^17^. While negative epistasis was pervasive, many deleterious mutations improved fitness in at least one alternative background, supporting the notion that epistasis expands the permissive portions of sequence space. Positive epistasis was rare, often long-range, and confined to a conformationally dynamic network of residues. Similarly, a comparison of DMS profiles in the PSD95 PDZ3 domain with two different ligands revealed positive epistasis in a set of adaptive positions, which belonged to a network of coevolving amino acids, termed a sector, that defines the constraints of PDZ ligand binding ^18, 19^. Epistatic and conditionally neutral mutations in a subset of adaptive positions distant to the ligand-binding site could mediate ligand class-bridging through allosteric ‘remodeling’ of the PDZ sector ^20, 21^. By providing an experimental means to link physicochemical variation at the amino acid level to epistatic phenomena at the protein level, deep mutational scanning led to new insight into the structural principles that underlie evolutionary adaptability.

DMS also suggested that epistatic interactions are enriched in mutation pairs that are close in structural distance. Comparable to using the co-evolution of amino acids to infer three-dimensional structure ^22–24^, epistatic interactions can be used as constraints for computational backbone structure determination ^25^. Similar to the idea of sectors that emerged from coevolution analysis, distinct clusters of structurally close residues with negative and positive epistasis were observed. While the former was related to protein stability, the latter was enriched for residues involved in ligand binding.

DMS clearly holds great value to protein science. Its value stems from the comprehensiveness of experimental datasets; comprehensiveness enables the development of quantitative models of the protein structure, function, and evolution. For single point mutations, this comprehensiveness is relatively easy to achieve, and the most common methods use a combination of degenerate oligos and ligation ^2–5, 7, 8, 17, 20, 26–28^ or error-prone PCR ^6, 9^. An alternative to degenerate oligos is programmed oligo pools ^29–31^ that can be used to encode specific codons, avoid stop codons, or target specific substitutions when constructing DMS libraries ^10, 13^. Because of the programmed nature of mutations, it is possible to detect and discard sequencing errors. Despite these advantages, programmed oligo pools have yet to be used for deep mutational scanning of double mutants. We recently developed Saturated Programmable Insertion Engineering (SPINE), which combines oligo library synthesis and multi-step Golden Gate cloning for programmed mutagenesis ^32^. Here we adapt SPINE as a programmable DMS technique. We validate this approach with an existing deep single mutational dataset in the PSD95 PDZ3 domain ^20^, and in addition, comprehensively characterize most double mutants. We corroborate earlier findings of wide-spread proximal negative epistasis and rare long-range positive epistasis in other position pairs for the PSD95 PDZ3 domain. Negative epistasis is enriched in the beta-sheets of the PDZ domain core where mutations likely exhausted threshold robustness ^14^. Positive epistasis is strongly enriched in pairs between sector ^19^ or conserved positions and residues that define the evolutionary clade of PDZ domains ^33^. Flex-ddG / Rosetta-Backrub-based simulations ^34^ suggest that positive epistasis has a structural mechanism in which a neutral mutation can compensate for the deleterious effect on protein stability of a second mutation. We find that conditional neutrality of mutations in these clade-specifying residues is required to compensate deleterious mutations in sector positions. This suggests that the specific epistasis between clade-specifying residues and the PDZ sector facilitated the evolutionary expansion and specialization of PDZ domains.

### SPINE mediated construction of comprehensive single and double mutant libraries

To construct mutant libraries, we adapted a method we recently developed for insertional mutagenesis that leverages programmable oligo library synthesis and multi-step golden gate cloning (**Fig. 1A**,, **Suppl. Fig. 1.1**) ^32^. Oligos were programmed to contain the desired mutational diversity in a custom algorithm (written for Python 3.7.3. and available at https://github.com/schmidt-lab/SPINE). To generate single mutant libraries, the wildtype PSD95 PDZ3 backbone ^20^ was used as the template, while double mutant libraries used the single mutation library as the target gene backbone template (**Fig. 1B**). This means that double mutants are always separated by a fragment boundary, which in our case means that they are at least 2 amino acids apart with an exponential increase in probability with greater distance from the fragment boundary (‘blackout regions’, **Supp. Fig. 3.2B**). All libraries at this step yielded greater than 100,000 colonies corresponding to greater than 30-fold coverage for single mutants and greater than 5,000,000 colonies corresponding to greater than 20-fold coverage for double mutants assuming 0.3% of the library has indels (the most common error with phosphoramidite chemistry ^35, 36^) and 15% of double mutations are in blackout regions. Due to inefficiencies of the DNA assembly, the wild-type original gene remained in the libraries at around 5% for the single mutation libraries (**Supp. Fig. 2.1B)** and 3.8% for the double mutation libraries (**Supp. Fig. 3.1B**).

**Figure 1.**
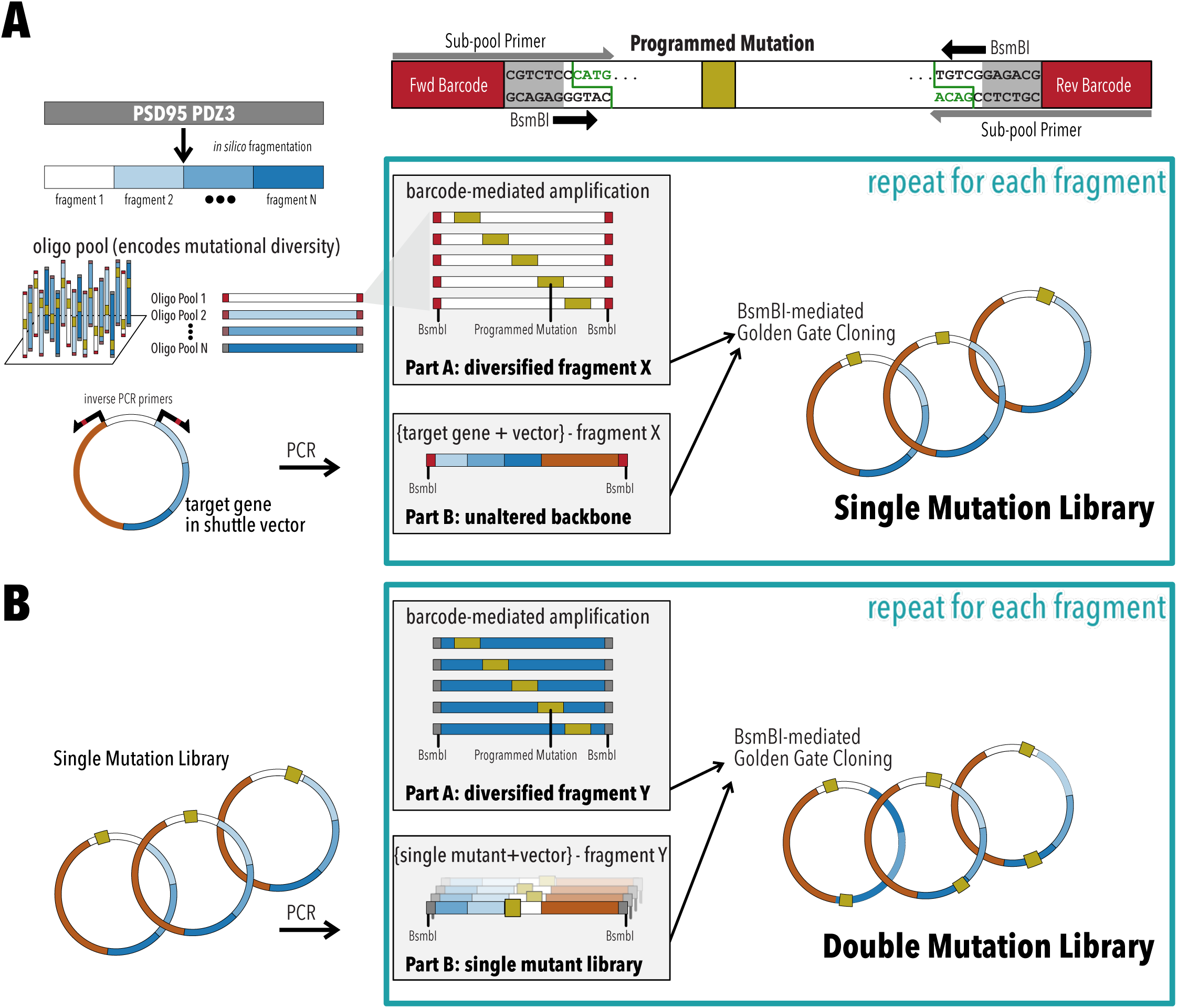
SPINE for comprehensive single and double mutant libraries. **A**, Single mutant libraries are constructed by dividing PSD95 PDZ3 into fragments and encoding the mutational diversity of each fragment within an oligonucleotide pool. Subpools (Part A), corresponding to each fragment can be amplified with subpool primers and combined with the corresponding unaltered backbone (Part B) by Golden Gate Cloning using BsmBI. Overhangs generated by this Type IIS restriction enzyme are unique for each fragment boundary. This process is repeated for each fragment to generate mutation sublibraries. Sublibraries are then combined into a complete Single Mutant library. **B**, Double Mutatant libraries are generated by the same process with Single Mutant libraries as input.

**Figure 2.**
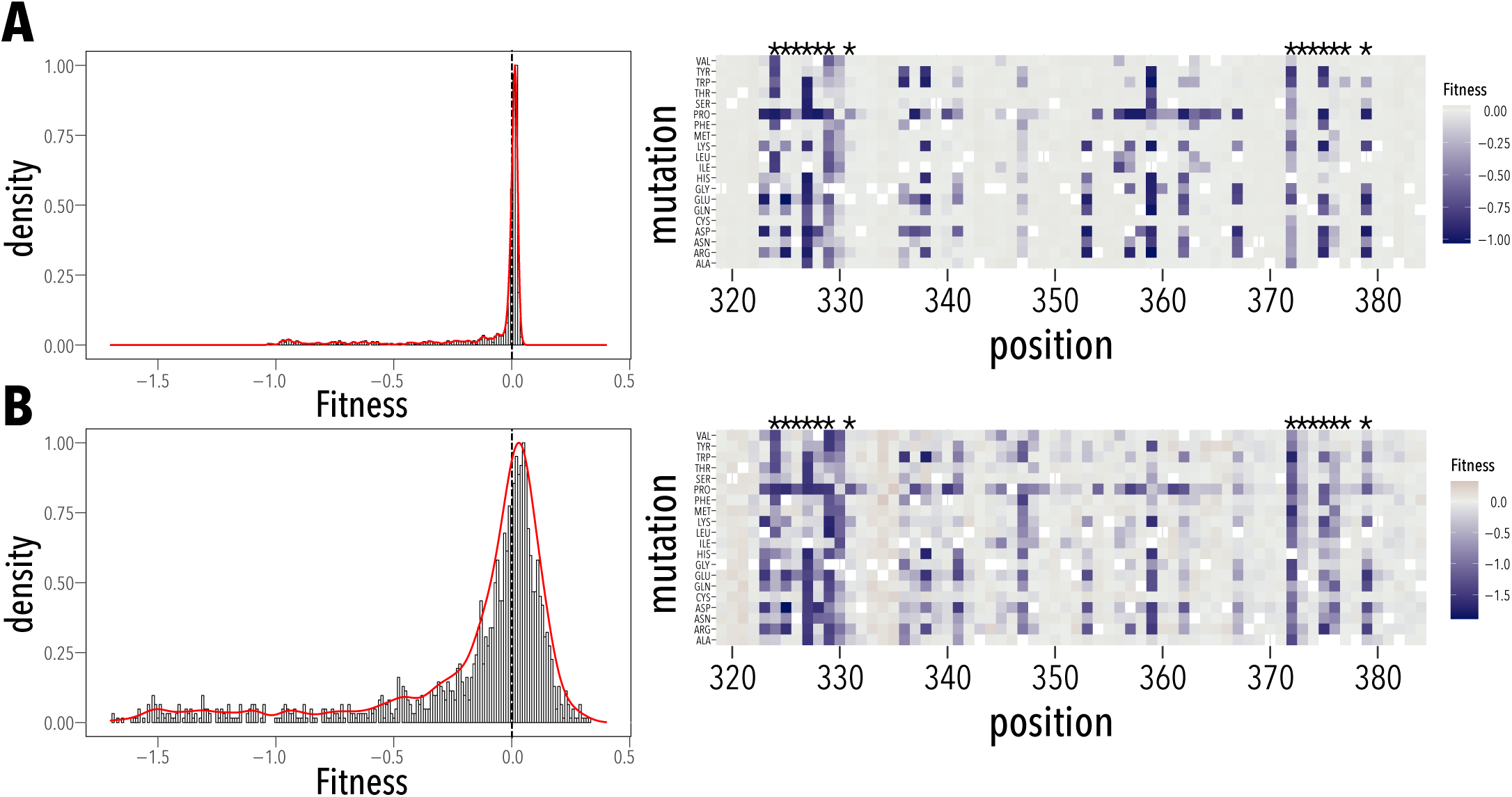
Single mutant fitness. **A**, Distribution of single mutant fitness (wildtype fitness = 0). While many single mutations in PSD95 PDZ3 are deleterious (fitness < 0) and few are beneficial (fitness > 0), most single mutants are neutral (fitness = 0; same as wildtype). Positional effect of each mutation is shown on the right. Asterisk (*) denotes residues in contact with the ligand. **B**, Distribution of single mutant fitness determined by McLaughlin et al. is in very good qualitative agreement, but has greater dynamic range.

**Figure 3.**
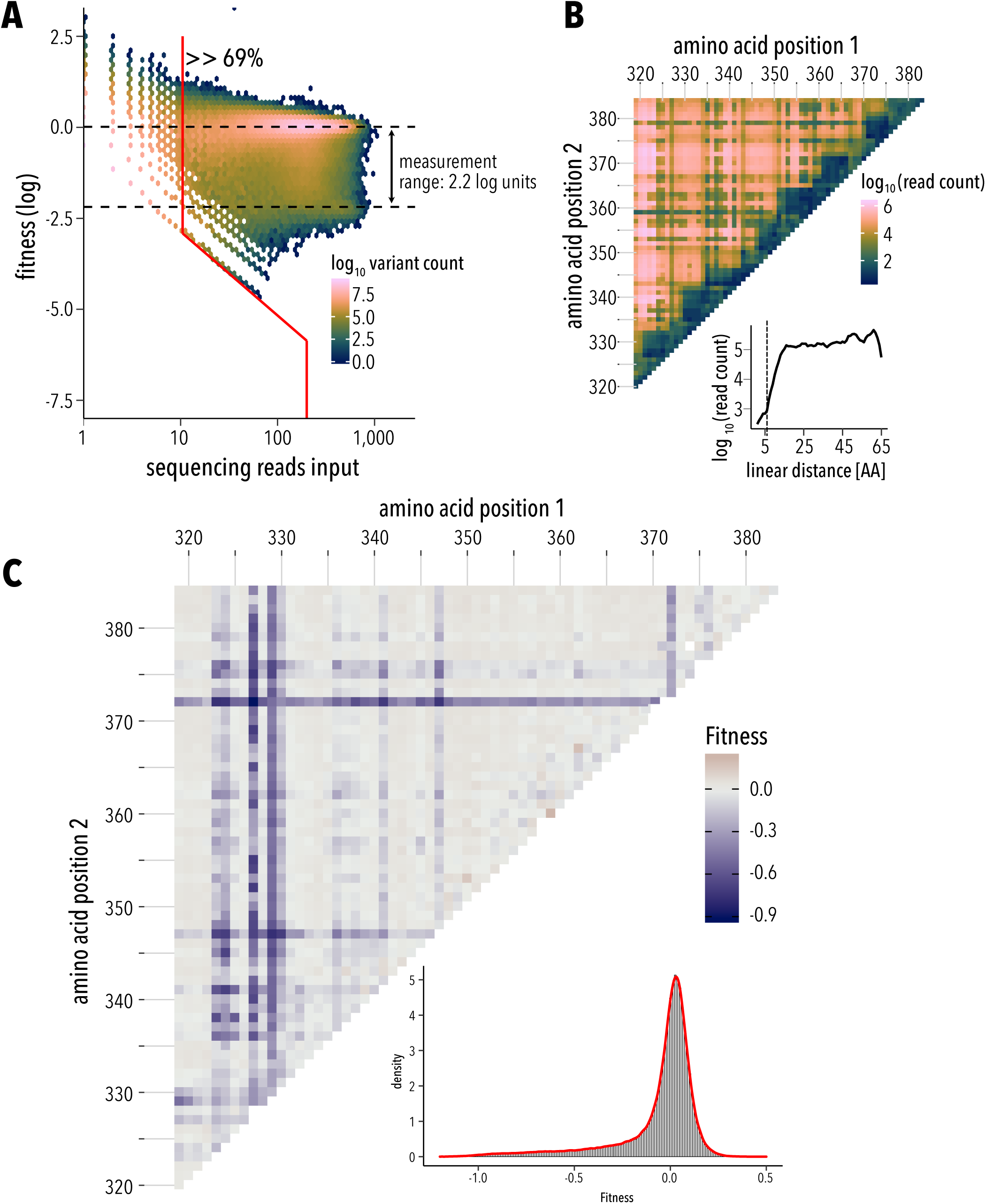
Double Mutant Fitness. **A**, Distribution of double mutant fitness by sequencing read counts. 69% percent of the 750,880 possible double mutants passed read quality threshold (200 reads, red line). Measurement range of the fitness assay is 2.2 log units (dashed horizontal lines, see Methods). **B**, Pairwise position map of double mutant shows that most missing mutants are close in linear sequence distance (<6 amino acids). **C**, Map of mean double mutant fitness averaged across all mutations for a position pair. Inset, double fitness distribution shows strong deleterious effects in many double mutants, but also improved fitness (compared to wildtype) for some.

### Single mutant library fitness

We assayed the effect of single and double mutants using an established bacterial two-hybrid system ^21, 37^ that couples the binding of PSD95 PDZ3’s ligand (CRIPT) to the expression of Chloramphenicol resistance (**Supp. Fig. 1.2**). We used NextGen sequencing to quantify the frequency of each mutant before and after antibiotic selection, and calculated each mutant’s relative fitness compared to WT:

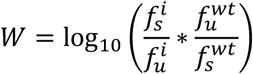

Count statistics showed that we have excellent depth for single mutant (greater than 100-fold at 95% of positions; median ∼6,500 counts, **Supp. Fig. 2.1A-B**). We determined fitness for all 1,235 single mutants, with similar replicates (R^2^ 0.93± 0.009) (**Supp. Fig. 2.1C, Supp. Fig. 2.2**). A median confidence interval relative to measured fitness for each single mutant (based on a 90% Poisson confidence interval) of 11.8% suggests good fitness measurement precision. Most single mutants are deleterious, while beneficial mutants are rare (**Fig. 2A**). Comparison of this deep single mutant dataset to earlier studies ^20^ showed good qualitative agreement (**Fig. 2B**), but we noticed that our dataset has less difference between the most beneficial and deleterious mutants. Furthermore, median fitness is not centered at 0 but shifted slightly to higher-than-wildtype fitness. A similar trend is in the reference dataset (**Fig. 2B**). This shift to higher fitness could be due to how the single mutant libraries are constructed. While each mutation in our approach was programmed to use a specific codon, McLaughlin et al. ^20^ used degenerate NNS primers, with N being any base and S being either G or C. This means that each amino acid substitution might be encoded by a different codon (e.g. Gly as either GGC or GGG) which are used at a different frequency in *E*.*coli* (15% and 37%, respectively). Codon content, in overexpressed proteins in bacteria, influences protein expression by affecting mRNA folding and translation, or overall cellular fitness ^38^. As expected from a programmable library generation method, empirical cumulative distribution functions for an NNS library and our library show that our approach used optimal codons more often (**Supp. Fig. 2.3A**). Better codon usage could result in slightly better expression and thus higher fitness in particular for neutral mutations. Comparing fitness effects of equivalent mutations in the McLaughlin et al. and our datasets, we find that there was a monotonic, but non-linear relationship between fitness for each mutant, with only a few (<10%) outlying residues (**Supp. Fig. 2.3B**). Outlying data points usually had lower confidence values, suggesting they are due to from limited sampling. Despite minor quantitative differences, the agreement of single mutation fitness validates our library construction method.

### Double mutant library fitness

Of the 750,880 possible double mutants, 648,138 (86%) are represented in the double mutant dataset, and 519,508 (69%) passed the read quality threshold with a median of 200 input reads for each position pair (**Fig. 3A, Supp. Fig. 3.2A**). Median fitness error relative to the measurement range is 0.15/2.2 log units = 6.9%, which is comparable to other double mutant datasets ^25^. Mapping read counts to linear distance in sequence reveals that most missing mutants are in close proximity (< 6 amino acids apart, **Fig. 3B, Supp. Fig. 3.2.B**). We expect this with our library generation technique in which two mutations never occur in the same fragment as only one mutation was encoded in each oligo (see methods). At 17-fold the median depth was lower than single mutants (**Supp. Fig. 3.1A-B**), however, replicates were in good agreement (**Supp. Fig. 3.1B**). Many double mutants have a strong deleterious effect on fitness, similar to single mutants (**Fig. 3C**), but improved fitness (compared to wildtype) is evident as well.

### Running median surface approach to calculating epistasis

If the relationship between measured fitness and underlying biophysical effects of mutations is non-linear, due to protein folding thermodynamics or cooperativity, a linear model of the fitness landscape will yield many non-specific epistatic interactions ^39^. To detect epistatic interactions that are specific, i.e. depend on identity of the involved residues and mutations, the global nonlinearity between biophysical effects of mutations and fitness phenotype must be estimated. A null-model to infer this landscape is a running median surface approach originally developed for determining protein structures from deep mutagenesis data ^25^. This approach also helps accounting for non-linearities that can result from varying uncertainty of fitness measurement (e.g. low read counts for low fitness variants), fitness measurements near the lower measurement limit of the fitness assay, and non-specific thermodynamic epistasis. We calculated epistasis using running quantile surfaces of average local fitness for double mutant data that was not impeded by measurement errors and passed read thresholds (15% and 44% of the double mutant space for positive and negative epistasis, respectively). A surface representing the average local fitness of double mutants is calculated using local polynomial regression (**Fig. 4A**). Then the 10^th^ and 90^th^ percentile fitness surface were calculated from a fitness distribution of a double mutant’s closest neighbors in single mutant space. Double mutants are categorized as positive epistatic if their surface-corrected fitness value was above the 10^th^ percentile, and negative epistatic it was below the 90^th^ percentile fitness surface (**Fig. 4B**). Overall, adding fitness of single mutants predicted double mutant fitness only moderately well (Spearman correlation coefficient 0.63, **Supp. Fig. 4.1A**) and many double mutants deviated from expected additivity, suggesting that epistasis is common in PSD95 PDZ3. Negative epistasis with an enrichment score > 2 or > 5 was observed in 72% or 16% of quantifiable position pairs, respectively (**Supp. Fig. 4.1B**). Conversely, positive epistasis enrichment greater > 2 or > 5 was found in 43% or 7% of quantifiable position pairs, respectively (**Supp. Fig. 4.1C**). Together this suggests that while epistasis is pervasive, weak negative epistasis is more common than strong positive epistasis.

**Figure 4.**
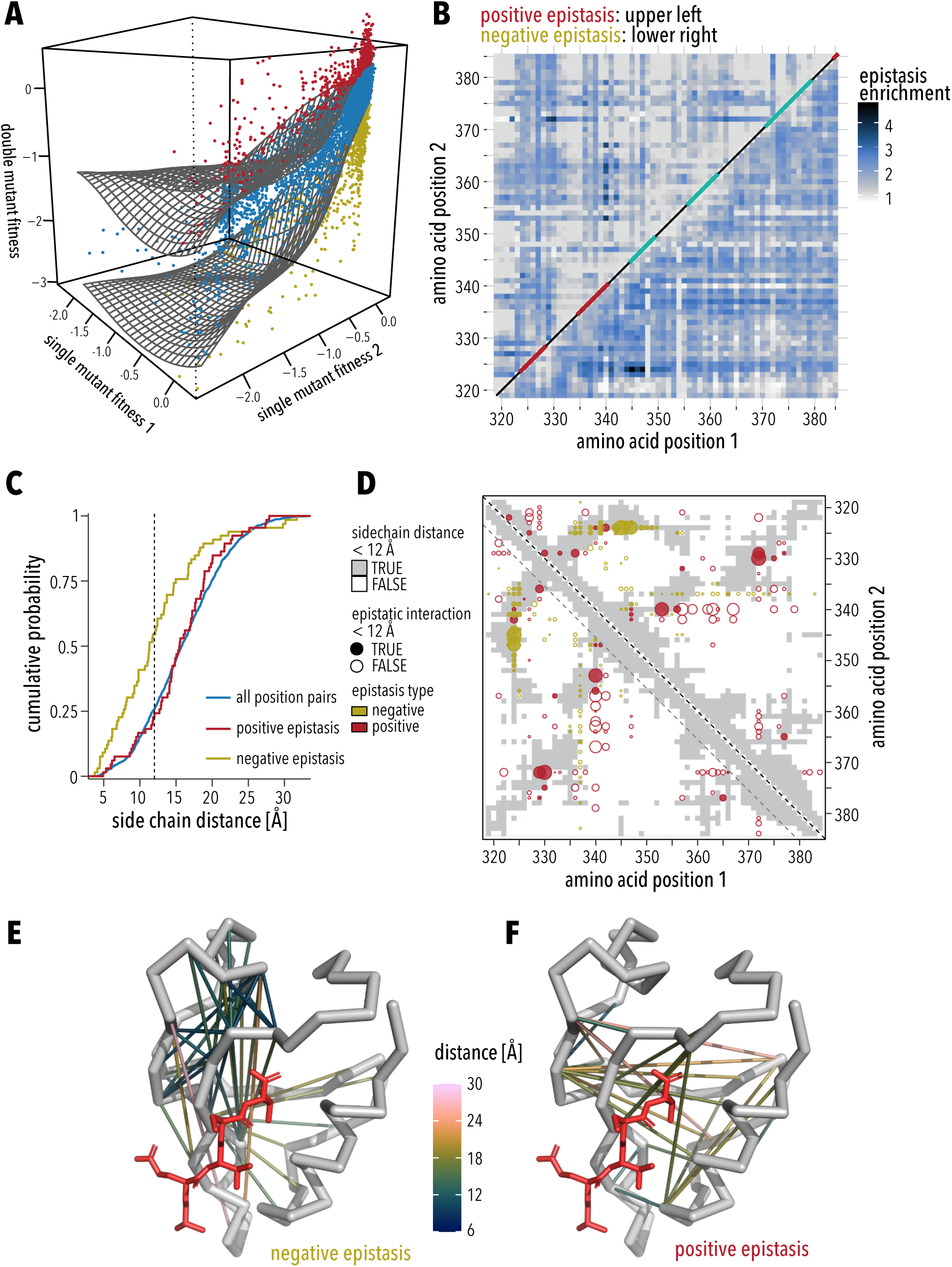
Running median surface approach to calculating epistasis. **A**, The fitness of each double mutant is along with the corresponding single mutant fitnesses. A surface representing the average local double mutant fitness is calculated using local polynomial regression. Double mutants are categorized as positive epistatic if their surface-corrected fitness value was above the 10th percentile (red dots), and negative epistatic it was below the 90th percentile (yellow dots). Blue dots represent non-epistatic double mutant pairs. **B**, Map of position pairs that are enriched in positive epistasis (upper left triangle), or negative epistasis (lower right triangle). **C**, Empirical cumulative distributions show that position pairs in PSD95 PDZ3 with negative epistatic interaction are more likely to be in proximity, while positive epistasis can occur over long distance. 50% of negative epistatic (yellow line) and 25% of positive epistatic pairs (red line) are <12Å apart, respectively. **D**, Position map that shows structural contacts (<12Å minimal side chain distance) as grey background. Epistasis enrichment is shown as dots, yellow for negative epistasis and red for positive epistasis. Dots for epistatic interactions between residues that form structural contacts are filled, those that are not in structural contact are empty. Magnitude of enrichment is indicated by dot size. Structure of PSD95 PDZ3 (PDB 1BE9) showing interacting residues pairs with negative epistasis (**E**) or positive epistasis (**F**). Connections between residues are colored by minimal sidechain distance. CRIPT ligand is shown in red.

### Spatial proximity of epistatic interactions

DMS in proteins ^2–4, 6–9, 13, 17, 26, 27^ and nucleic acids ^40–43^ have suggested that epistasis is more likely to occur between proximal residues as opposed to distal residues. This is the basis of structure prediction from DMS data, which has been demonstrated for several model proteins ^22,25^. Comparing distance distributions in PSD95 PDZ3 shows that position pairs with epistatic interactions are more likely in proximal pairs (<12A minimal side-chain heavy atom distance, scHAmin, **Fig. 4C**). This trend was mostly driven by negative epistatic position pairs in that 50% of negative epistatic and 25% of positive epistatic pairs are <12A apart. While a small cluster of proximal pairs (5-7A) with positive epistatic interaction can be seen in the data, most appear to be distal interactions (>12A). Note that missing data is unlikely to affect to this observation as only 4% of residues pairs with a linear sequence distance of < 6 amino acids have a minimal side-chain distance of > 12A. The distinction between proximal negative epistasis and distal positive epistasis is apparent when we overlay the type and magnitude of epistasis onto the PSD95 PDZ3 contact map (**Fig. 4D**) or structure (**Fig. 4E-F**). While the position pairs with enriched negative epistasis make structural contacts (filled yellow circles on grey background), this is not the case for positions with enriched positive epistasis (open red circles on white background), which often occurs over long distances. Protein folding is mediated by structural contacts, for example hydrophobic interactions in the core of the protein ^44–46^. This explains why fitness of double mutants is particularly impaired when both positions are mutated to disruptive (proline), bulky (tryptophan), or charged (glutamate, aspartate) amino acids (**Supp. Fig. 4.2A**). Grouping double mutant fitness by descriptors that capture amino acid property of wildtype and mutants illustrates this trend further. Fitness is strongly decreased in double mutants if both wildtype positions are aromatic or non-polar (**Supp. Fig. 4.2C**). The stratification of epistasis in double mutants by amino acid paints a different picture (**Supp. Fig. 4.2B**&**D**). As expected, mutations to bulky aromatics (Phe or Trp) or proline show strong negative epistasis in the background of proline and tryptophan mutations at a second site. In the background of second site proline or tryptophan mutations, negative epistasis is also observed for many charged and polar mutations. The same charged or polar mutation in the background of small non-polar (valine, leucine, isoleucine) mutations, however, show positive epistasis (**Supp. Fig. 4.2B**). Sign dependence of epistasis on background mutation type is strongest when aromatic residues are mutated to charged residues (**Supp. Fig. 4.2D**). Together this data suggests a multi-faceted mechanism for how epistasis arises in PSD95 PDZ3.

### Strong negative epistasis arises from exhausted threshold robustness

Theoretical and experimental work supports a mechanistic connection between negative epistasis and threshold robustness ^14, 47–49^. Single mutations may have little impact on fitness if their effect is buffered by excess stability. If the first mutation largely exhausts this stability threshold, subsequent mutations will have a non-additive (i.e., epistatic) impact on fitness even if individually they minimally impact fitness. 2D histograms of the individual fitness of single mutations binned by epistasis provides a way to visualize that exhausted threshold stability can explain strong negative epistasis in PSD95 PDZ3. For the least fit double mutant position pairs (2.3 percentile), negative epistasis was common (mean epistasis score = -1.39± 0.008, **Fig. 5A**). Epistasis was most negative when single mutants were neutral, i.e. individually had minimal impact on fitness (**Fig. 5B**). This suggests that single mutants already exhausted excess stability or ligand binding activity such that a second neutral mutation led to a strong decline in fitness. Conversely, in double mutants that had near wildtype fitness or even better than wildtype fitness (97.7 percentile) positive epistasis was prevalent (mean epistasis score = 0.5±0.001, **Fig. 5A**). For this group, positive epistasis was strongest when a deleterious mutation occurred in the background of a neutral mutation (**Fig. 5C**).

**Figure 5.**
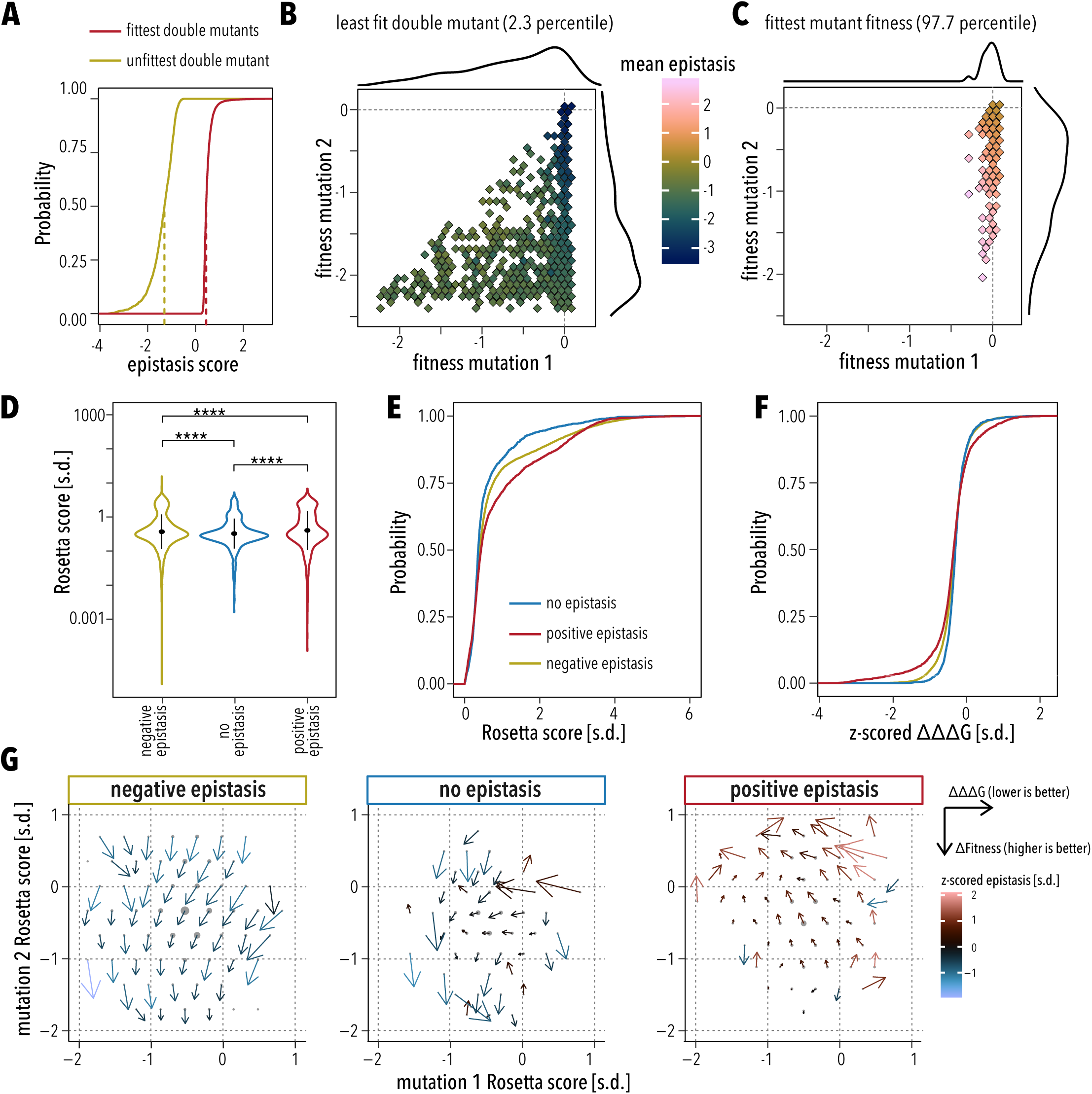
Strong negative epistasis arises from exhausted threshold robustness. **A**, Empirical cumulative distribution function of epistasis scores in the least fit (yellow line) and fittest (red line) double mutants. Vertical dashed line indicate median epistasis for each double mutant set. **B-C**, Binned scatterplot of single mutant fitnesses for the least fit (**B**) and the fittest (**C**) double mutants. Fill color indicates mean epistasis of double mutants in each bin. The number of double mutants represented by each bin is indicated as marginal density plots. For the least fit double mutants, epistasis was most negative when mutation 1 was neutral, suggesting that this mutation already exhausted excess stability of the protein. For the fittest double mutants, positive epistasis was strongest when a deleterious mutation 2 occurred in the background of a neutral mutation 1. **D**, Rosetta energy scores from flexddG backrub calculations of double mutants in positions pairs enriched for negative epistasis (yellow), no epistasis (blue), or positive epistasis (red). Black dot indicates mean, vertical black line indicates standard error. Difference between means was compared by two-sided Wilcoxon test;**** indicates p-values < 0.00001. **E**, Empirical cumulative distribution of Rosetta energy scores from (**D**). **F**, Empirical cumulative distribution function of ΔΔΔG, the calculated difference in protein stability between a double mutant and summed stability of respective single mutants. Lines are color-coded as in (**E**). **G**, Binned quiver plots of Rosetta energy scores for single mutations distribute similarly in each category (grey dots). Dots size indicates the number double mutant represented in each bin. Arrow direction and length indicates sign and magnitude of ΔΔΔG (additional stabilization in the double mutant compared to summed single mutants) and Δfitness (additional fitness in the double mutant compared to summed single mutants). Arrow color indicates mean epistasis.

### Residues for which double mutations improved protein stability are enriched for positive epistasis

To investigate the mechanistic link between fitness and epistasis we used the “flex ddG” protocol ^34^, implemented in Rosetta, to model the effect of independent and pairwise mutations in PSD95 PDZ3 on protein stability. This protocol first generates conformational ensembles by a local sampling of backbone and side-chain flexibility using Rosetta’s backrub algorithm. After repacking and global minimization, changes in folding free energy are estimated between the simulated wildtype protein vs. a single or double mutant (ΔΔG). Overall, there was a weak correlation between fitness and estimated ΔΔG (R = -0.25, p < 2.2e-16) and no correlation between epistasis and estimated ΔΔG (R = -0.073, p<2.2e-16). However, mutations in residues that are enriched for either negative or positive epistasis are more destabilizing (larger ΔΔG) than mutations in residues pairs without epistasis (null set, **Fig. 5D-E**). We then calculated the difference in protein stability between a double mutant and each respective single mutant:

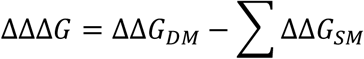

A negative ΔΔΔG indicates that the double mutant is more stable than the added independent effects of single mutants. Inspection of the empirical cumulative distribution function for ΔΔΔG revealed that mutations in residue pairs enriched for positive epistasis are more likely to result in greater protein stability than expected from the added effects from each single mutant (t.test p< 0.0001, **Fig. 5F**). No stabilizing effect is observed between residues pairs that are enriched in negative epistasis.

What is the relationship between epistatic stabilization (ΔΔΔG, lower is more stable) and non-additive fitness (Δfitness, higher is better)? Reiterating the weak or absent correlation between fitness or epistasis with calculated protein stability, we find a similar range and distribution of z-scored Rosetta scores for single mutants in negative epistasis, no epistasis, and positive epistasis subsets (**Supp. Fig. 5.1**). However, when we use a vector representation to overlay ΔΔΔG and Δfitness (**Fig. 5G**, arrows) onto single mutant Rosetta scores (**Fig. 5F**, bin centers represented as grey dots), we observe distinct differences between negative and positive epistasis. In position pairs that are enriched for negative epistasis, the arrows generally point straight down. This means that there generally is little additional stabilization in the double mutant (ΔΔΔG ∼0) and that double mutants are less fit than predicted from summed single mutant fitness. In position pairs that are enriched for positive epistasis, however, arrows generally point to the left and up. This means double mutants are generally more stable than predicted from the protein stability of single mutants, and that the fitness of double mutants is greater than predicted from the fitness of single mutants. This effect was strongest in position pairs that had the highest enrichment of positive epistasis (**Fig. 5G**, right panel, arrow color). In aggregate this suggests a mechanism for the positive epistasis observed in these residue pairs: mutations that in the wildtype PSD95 PDZ3 background would be destabilizing are less stabilizing in the background of a second mutation, which itself is neutral (**Fig. 5C**) and does not alter stability (ΔΔG ∼0, **Fig. 5G**).

### Epistasis and PDZ protein sectors

The premise for 3D structure prediction from deep mutational scanning is that specific epistasis is enriched between proximal residues and is less common between distal residues ^22, 25^. While residues pairs with enriched negative epistasis follow this trend in our dataset, positive epistasis more frequently occurs over longer distances (**Fig. 4C-F**). We therefore sought other features of PSD95 PDZ3 that could explain the observed patterns of positive epistasis (**Supp. Fig. 6.1A**). As the first feature, we calculated positional conservation using the Kullback-Leibler divergence of positional amino acid frequency in a PDZ family alignment ^33^ versus the amino acid frequency in vertebrate protein deposited in Uniprot. The second feature is based on previous DMS in PSD95 PDZ3 that defined positions that show epistasis with respect to binding wildtype CRIPT ligand vs. a class-switching T_-2_F ligand ^20^. The third feature is based on a reanalysis of that dataset, to define a set of adaptive positions that are either class switching (gain of binding to T_-2_F with loss of binding to CRIPT) or class-bridging mutations (gain of binding to T_-2_F and maintain binding to CRIPT) ^21^. The fourth feature describes a residue’s spatial proximity to the ligand ^21^. The fifth feature is based on studies in PSD95 PDZ3 that proposed sparse networks of co-evolving residues, ‘sectors’ ^18, 19^, as the mechanistic basis for a protein’s function. Sector positions are sensitive to mutations whereas non-sector positions are more tolerant, which suggested that the sector architecture provides mutational robustness and adaptability ^21^. The sixth feature is evolutionary sequence conservation (coupling) among sets of residues, which can point to an interdependence of phenotypes that arise from genetic variation ^23^. We tested which feature can explain positive epistasis using Fisher’s Exact Test, with the null hypothesis of independence. Positive epistasis (>3sd, **Supp. Fig. 6.1C**) was enriched in conserved residues (p-value 0.002), in positions that enable ligand class-switching and class-bridging (p-value 0.03), strongly in positions that contribute to ligand specificity (p-value 2.5×10^−6^), and in sector positions (p-value 8.5×10^−5^). Positive epistasis was not enriched in residues that contact the ligand (p-value 0.24) nor in evolutionarily coupled positions (p-value 0.76). For negative epistasis (>2sd) the null hypothesis was not rejected for any category (p-value > 0.05, **Supp. Fig. 6.1B**), suggesting that perhaps it is determined by perhaps other properties, such as protein stability and folding. This is in line with our observation that negative epistasis occurred predominantly along core beta-sheets (**Fig. 4E**). In aggregate, these results reaffirm the connection between epistasis and evolutionary processes such as adaptation ^14, 16^. They provide further support for the theory that protein sectors originate from non-local (i.e. long-range, allosteric) interactions between residues that provide conditionally neutral capacity –here measured as positive epistasis– to adapt to fluctuating selection pressures and fitness conditions ^20, 21^.

**Figure 6.**
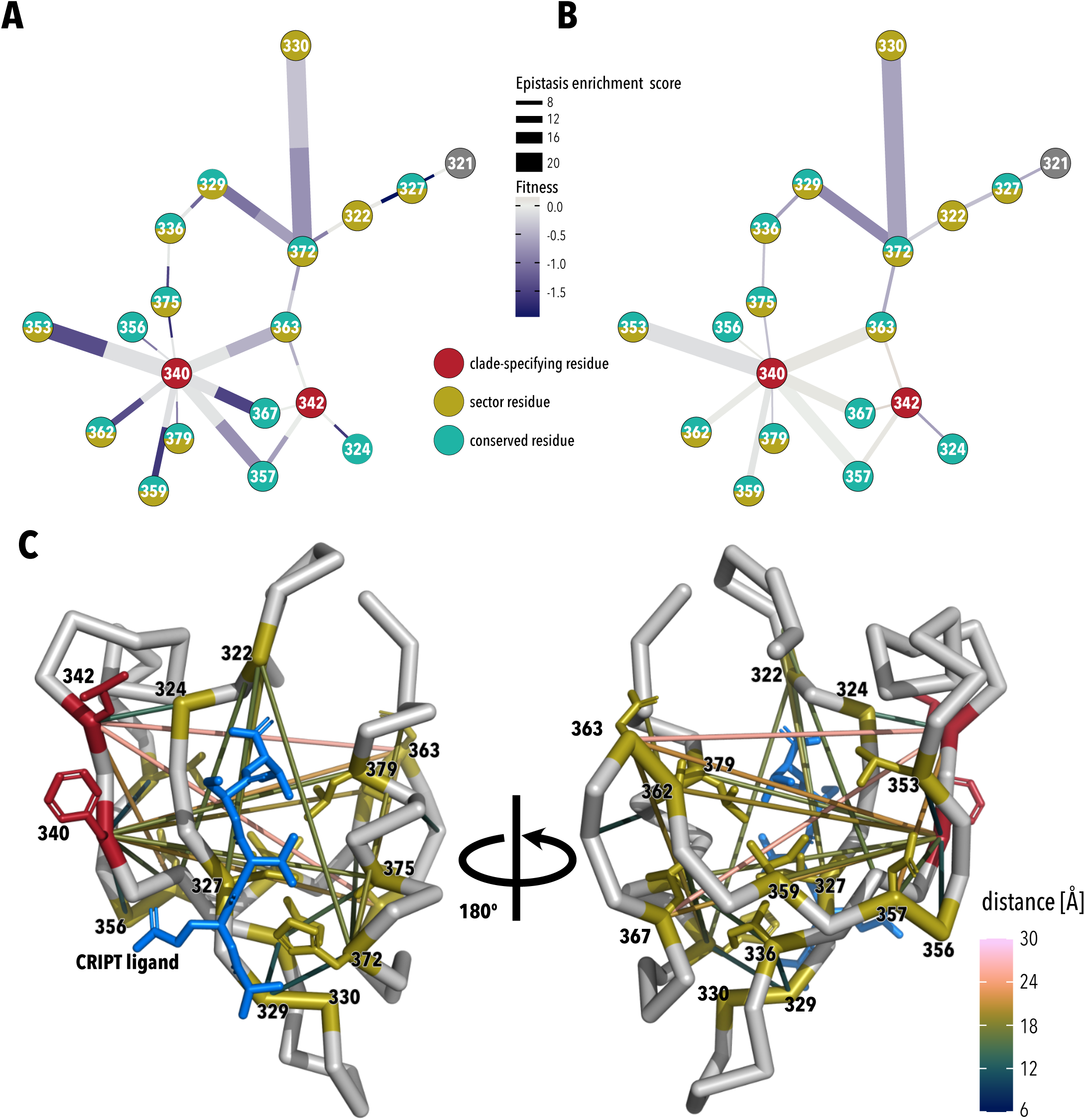
Positive epistasis in clade-speciic positions. **A**, Network diagram of amino acid positions with an positive epistasis enrichment score 3. Nodes are colored by category: clade-specifying residue (red), sector residue (yellow), or conserved residue (teal). Edge thickness between nodes indicates magnitude of epistasis. Edge are divided into two sections; color of the section adjacent to a node indicates median fitness of the node’s single mutants. With the exception of two clade-specifying residues, almost all epistatic interactions are mediated by sector and/or conserved residues. Single mutations in clade-specifying positions are neutral, while mutations in sector and conserved positions are deleterious. B, Same network diagram as in (A), but now edge color indicates fitness of the double mutant. In the background neutral mutations in positions 340 and 342, median fitness of otherwise deleterious second site mutations was improved, while double mutant in residue pairs that did not involve clade-specifying residues were still deleterious. C, Structural mapping of clade-specifying (red) and sector and/or conserved positions (yellow) with strong epistasis. Color of each connections indicates side chain distance. F320 and L342 are not in direct contact with the CRIPT ligand (blue).

### Positive epistasis in clade-specific positions

The special relevance of epistasis in PDZ family diversification becomes even more evident from a network analysis of positive epistatic interactions in PSD95 PDZ3. It reiterates that almost all strong interactions (enrichment score > 3sd) are mediated by sector and/or conserved residues (**Fig. 6A-B**, yellow and blue circles). The two exceptions are positions F340 and L342 (red circles), which do not belong to the PDZ sector and are not evolutionary coupled with other PDZ residues, but clearly form the central hubs of a network from which interactions with evolutionarily-coupled residues radiate. Another smaller hub is centered around H372, which is important for ligand class-switching ^20, 21^. This organization around F340 and L342 is noteworthy as they belong to a group of residues that identify the clade of PDZ domains. PDZ domain usage expanded greatly along the stem leading from choanoflagellates (the closest living relatives of animals), and later metazoans. A comparison of global entropy vs. within-clade entropy of all positions revealed that six residues (F340, I328, D332, G333, S339, L342 in PSD95 PDZ3) alone can classify >95% of PDZ domains to the correct evolutionary lineage ^33^. Two of these classifying residues (D332 and G333) are located in a loop with frequent deletions and insertions in PDZ domains. Two other classifying residues are in direct contact with the ligand (I328 and S339 in PSD95 PDZ3) and have negative epistasis in our dataset. F340 and L342, which are strongly enriched for positive epistasis, do not form direct contact with the ligand (**Fig. 6C**). Median fitness across all single mutants in F340 and L342 is near wildtype, a near-neutral phenotype, while single mutations in connected sector and/or conserved residues decreased median fitness (**Fig. 6A**). In the background of neutral mutations in F340 or L342, however, median fitness upon mutating these connected residues was rescued or slightly improved over wildtype (**Fig. 6B**). This data argues that at least two of the six PDZ clade-specifying residues are intimately connected to a function-defining coevolving set of amino acids. The fact that F340 and L342, unlike their coevolving interaction partners, have remained unchanged over the course of 600 million years of animal evolution suggests a key role for long-range epistatic interactions between clade-defining and function-defining residues in not only in PDZ expansion and specialization, but also maintenance of ligand specificity.

## Discussion

Deep mutational scanning is an important tool to study epistasis in proteins. Comprehensively measuring the effects of mutations is key to map protein fitness, at least in the local sequence neighborhood, with high resolution. The underlying mutant libraries are commonly generated through a combination of degenerate oligos (encoding mutational diversity as NNS or NNK codons) and ligation, or an error-prone PCR process. Recently, programmed oligo pools have found wider adoption as an economical alternative to produce oligos carrying specific substitutions, which makes it easier to detect sequencing errors. Oligo pools, to our knowledge, have not been used to generate large scale double mutant libraries, which prompted us to adapt Saturated Programmable Insertion Engineering (SPINE) for this application. Compared to error-prone PCR, which is easier to implement, SPINE has the advantage of stringent control the sequence, location, and number of mutations. Compared to degenerate oligo library design, e.g. used by Olson et al. ^17^, SPINE’s main advantage lies in its unambiguous assignment of sequencing read to mutations. Because mutational diversity is encoded as specific codons (instead of degenerate codons), we do not need internal barcodes to remove sequencing or oligo synthesis errors. Furthermore, SPINE uses 4-bp overhangs for Golden Gate assembly that uniquely define each fragment boundary as opposed to the degenerate K/M scheme. This means that the entire library can be assembled in a single reaction because each mutagenized fragment only ligates to the specific backbone amplicon that is missing this fragment, which simplifies library construction workflows. The downside of this approach is that two mutants must be at least 2 amino acids apart and have there is a lower probability of observing double mutants separated by less than 6 amino acid. (**Supp. Fig. 3.2B**). Double mutant libraries constructed with SPINE therefore contain ‘black-out’ regions with low coverage. Given the relative equivalence to degenerate oligo-based library construction, what benefit does SPINE offer? One potential benefit relates to the question of how epistasis affects long-term evolution of proteins, which requires investigation of higher-order interactions and epistasis. Experimental access to these experiments is readily achieved with SPINE. Any number of fragments, representing specific regions of a protein and each containing every single site mutation, can be assembled, in a single reaction, according to the logic encoded in the 4-bp overhangs. Because SPINE requires no error correction to distinguish mutations from sequencing or oligo synthesis errors, it makes more efficient use of sequencing platform throughput.

In agreement with other studies, we found that weak epistasis was prevalent while strong epistasis was rare. Negative epistasis was enriched in position pairs that make structural contacts, suggesting that one underlying mechanism is direct interaction. A similar enrichment of epistasis which is specific (i.e., described not only by effect size but also mutation identity) in proximal residues was observed in the analysis of the GB1 double mutant dataset and this formed the premise for the 3D structure prediction from deep mutagenesis data ^22, 25^. Specific epistasis is thought to leave a strong signal in the co-evolution of directly interacting residues ^16^. Statistical models that use a maximum-entropy approach to identify co-evolution in natural sequences perform better when interactions between all residue pairs in a protein are explicitly modeled to account for epistasis, and these models particularly improve predictions involving sets of proximal residues ^23^. Despite enrichment, our data, in particular for positive epistasis (**Fig. 4C, D, E**), and other studies show that epistasis is not exclusive to structural contacts ^17, 22, 25^. This suggests epistasis can occurs through a mechanism other than direct contact.

For PSD95 PDZ3, cooperative changes in sparse networks of residues (protein sectors ^18,19^) may explain such indirect effects of long-range epistatic interactions. By assessing the impact of a global single mutations on PDZ binding the native CRIPT ligand or the non-native T_-2_F ligand, statistically significant epistasis was observed in a set of residues that largely overlapped with the PDZ protein sector ^20^. For four residues (G322, G329, G330, and H372) positive epistasis was so strong that certain mutants at these positions were class-bridging or class-switching with respect to T_-2_F binding. Only H372 is in direct contact with the ligand suggesting that mutational effects in the protein sector mediated epistatic effects on ligand binding. The structural basis for this was described in a later study ^21^. Conditionally neutral (adaptive) mutations in sector positions, for example in G330, stabilized additional conformational states to enable ligand class-bridging, which was subsequently exploited by mutations in H372 for class-switching. Neutral G300 mutations are therefore crucial for the adaption of PDZ to bind new ligands. Consistent with these studies, we recorded the strongest positive epistasis signal between H372 and G329 or G330 (**Fig. 4B**, upper left triangle) and we could establish a relationship between positive epistasis, adaptive mutations, and sector positions (**Supp. Fig. 6.1B**). In fact, co-evolving residues clearly organize into a network that is strongly enriched for positive epistasis (**Fig. 6**).

Two residues (F340 and L342) are part of this positive-epistasis network and have strong epistatic interactions with sector and/or conserved positions but are themselves not co-evolving with other PDZ residues nor mediating adaptation to new ligands. A phylogenetic analysis of the major clades of bilaterian PDZ domains revealed that these residues are not conserved across the PDZ family. They are, however, highly conserved within each PDZ clade ^33^. In 600 million years of animal evolution, over which the PDZ family saw drastic evolutionary expansion and gained more than 300 PDZ domains, these positions have remained constant. This aligns well with the evidence that positions with strong epistasis have a low likelihood of reversion due to acclimatization ^50^. In light of apparently strong purifying selection, the epistatic interaction of F340 and L342 with sector positions in PDZ suggests a mechanism for how clade-specifying residues may have aided the evolutionary adaptation to different PDZ ligands. Restricted and rugged fitness landscapes due to negative epistasis constrict evolutionary pathways, while positive epistasis can provide paths that would otherwise be blocked by deleterious mutations and thus accept a wider range of mutations ^16, 39^. Conditionally neutral mutations in positions 340 and 342, through non-local allosteric mechanisms, stabilize the otherwise deleterious effects of adaptive mutations in sector positions, which by its cooperative nature, affects ligand binding. In some cases, this results in gain of function for new ligands, and if new ligand specificity provides a selective advantage these mutations become fixed. Positions 340 and 342 are then part of the genetic background that determines ligand specificity. Because subsequent mutations in these positions would negate their stabilizing effect and compromise ligand specificity, positions 340 and 342 now have come under purifying selection and thus emerge as clade-specifying residues. Future studies that assess specificity of PSD95 PDZ3 single and double mutants towards members of a randomized peptide ligand library are needed to test whether this adaptive path involving mutations in position 340 and 342 and sector positions is plausible.

Based on mutagenesis in PDZ and other proteins ^49, 51–53^, an ‘outside-in’ principle for protein adaption was proposed, in which adaption begins with mutations distant from active sites. Distant mutations are often neutral because their spatial separation from active sites makes it less likely that they break existing function. At the same time, distant mutations could provide access to new conformational states that are exploited by mutations closer to the active site. In the limit that PDZ is a small protein, the greater spatial separation of F340 and L342 from the ligand binding site, compared to sector positions (**Fig. 6C**), may be significant in light of this theory. The data presented here and previous work ^20, 21^ are consistent with the idea that residues in spatial proximity to the ligand (**Fig. 2**, asterisks) are the primary determinants of ligand binding. Adaptation to new ligands involves mutations in sector positions that are typically several shells away from the binding pocket. The effect of sector mutations is modulated by even more distant residues through positive epistasis. According to this model, an outside-in hierarchy of layers (clade-specifier > sector > active site) act in concert to define binding phenotype. Further experiments are needed to rigorously test this idea and generalize it to other proteins, but extensive biochemical data and sector descriptions are available for kinases ^54^, dihydrofolate reductase ^53^ and cryptochrome ^55^ whose functions are compatible with a DMS-style fitness assay. SPINE could help construct the required large-scale double and higher-order mutant libraries.

## Materials and Methods

### Oligo design

Oligo sequences are generated using a custom algorithm (written for Python 3.7.3. and available at https://github.com/schmidt-lab/SPINE).

#### Target gene fragmentation

The PSD95 PDZ3 gene was a gift from Rama Ranganathan. The PDZ sequence was replaced with a few alternative codons to remove recognition sequences for the restriction enzymes used in cloning. This new sequence was synthesized by Genscript before sequencing the donated plasmids. The PDZ sequence was divided into 10 evenly distributed fragments to the nearest codon (**Fig. 1A, Suppl. Fig. 1.1A**). Each fragment break site is adjusted to create unique cut site overhangs for Golden Gate cloning. If adjusting one fragment position causes any fragment to exceed the maximal length, the other fragments are adjusted to equalize fragment distribution below this length threshold (**Suppl. Fig. 1.1B)**.

#### Target gene primer design for inverse PCR

Forward and reverse plasmid primers are designed to amplify the backbone for each target gene fragment (**Supp. Fig. 1.1B**). Additional non-annealing sequences are added to the primer’s 5’ end encoding for inward-facing BsmBI recognition sites with the cut site including the first and last codon of the fragment (three bases) plus one base extension for the four base cut site. These primers are optimized for melting temperature and specificity by adjusting the length of the 3’ end. Melting temperatures are set between 55°C and 61°C based on calculations from both Sugimoto *et al*. ^56^ and SantaLucia and Hicks ^57^. A primer is flagged as nonspecific if annealing temperatures are greater than 35°C at any other position in the plasmid. Non-specific primers are made specific by extending the primer or, if max melting temperatures are exceeded, the fragmented site is adjusted.

#### Design oligos that encode each mutation

For each gene fragment, a loop is run to generate oligos for 19 mutations for each position within that fragment, starting after the first codon and ending before the last codon to account for the restriction enzyme cut sites. Therefore, to account for the cut sites, sequential fragments overlap by two codons. Mutations were generated by selecting each of the 19 amino acid codons weighted by their codon usage frequency in *E. coli* (obtained from Genscript) (**Suppl. Fig. 1.1C**). Codon usage frequencies below 0.1 were removed before selection with bias. The selected mutant codon replaced the existing wild type codon when assembling the oligo. Oligos consist of a bio-orthogonal barcode for specific subpool amplification, BsmBI recognition sites, and the fragment sequence with a mutation (Figure 1B). Barcodes are courtesy of the Elledge lab ^58^. In detail, each oligo starts with a forward subpool specific barcode, appended with a forward-facing BsmBI recognition sequence plus one base to bring the cut site into frame. The fragment with a mutation is appended followed by one base to bring the cut site into frame, a reverse facing BsmBI sequence, and a reverse subpool specific barcode. Due to the inefficiencies of the DNA assembly, the wild-type original gene remains in the libraries at around 5% for the single mutation libraries and 1.5% for the double mutation libraries, which serves as an internal control.

#### Design of subpool amplifying oligos

Forward and reverse subpool specific oligo primers are generated by testing annealing of a candidate primer sequence to the respective barcode, BsmBI recognition, and cut sequence. These primers are optimized for annealing temperature as described above, however, because the 3’ end is limited to the cut site, melting temperatures are optimized by adjusting the 5’ end or swapping the barcode sequence (**Suppl. Fig. 1.1D**).

#### In silico *quality control*

A final *in silico* quality control is run to check for the creation of new BsaI or BsmBI recognition sites and check for nonspecific subpool primers across all oligos (**Suppl. Fig. 1.1E**). If a BsaI or BsmBI recognition site is created, a codon within that recognition site will be changed to an alternative codon maintaining the amino acid sequence. Non-specific subpool primers are identified by an annealing temperature greater than 35°C for any position in any oligo other than the designed position. If a primer is non-specific, that subpool amplification barcode is replaced with another barcode and quality control is repeated. All oligos and primers are exported as FASTA files for ordering.

### Oligo library subpool amplification

A 7.5K oligo library synthesis (OLS) pool containing 1577 oligos for the PSD95 PDZ3 gene. OLS subpools corresponding to a given gene fragment were PCR amplified using Primestar GXL DNA polymerase (Takara Bio) according to the manufacturer’s instructions in 50 µl reactions using 1 µl of the OLS pool as the template and 25 cycles of PCR. The entire PCR reaction is run on 1% agarose gel and the DNA at 230bp was purified (Zymo Research). See Supplemental Figure S2.

### Assembly of single mutation OLS fragments and target gene backbone

To insert the OLS subpools into target gene backbones, complementary BsmBI sites to those on the OLS fragments of a respective subpool were added by PCR using Primerstar GXL DNA polymerase (Takara) and 100 pg of wildtype channel as template DNA (Supplemental Figure S3A). PCR products were purified using a 1% agarose gel to remove any undesired PCR by-products.

Target gene backbone PCR product with added BsmBI sites and the corresponding OLS subpools were assembled using BsmBI-mediated Golden Gate cloning ^59^ (Supplemental Figure S3B). Each 20 µl Golden Gate reaction was composed of 100 ng of backbone DNA, 20 ng of OLS subpool DNA, 0.2 µl BsmBI (New England Biolabs), 0.4 µl T4 DNA ligase (New England Biolabs), 2 µl T4 DNA ligase buffer and 2 µl 10 mg/ml BSA (New England Biolabs). These reactions were placed in a thermocycler with following program: (i) 5 min at 42°C, (ii) 10 min at 16°C, (iii) repeat (i) and (ii) 40 times, (iv) 42°C for 20 min, (v) 80°C for 10 min. Reactions were cleaned up using Zymo Research Clean and Concentrate kits, eluted in 10 µl of elution buffer, transformed into E. cloni®10G chemically competent cells (Lucigen) according to manufacturer’s instructions. Cells were grown overnight at 30°C to avoid overgrowth in 50 ml LB with 40 µg/ml kanamycin with shaking, and library DNA was isolated by miniprep (Zymo Research). A small subset of the transformed cells was plated at varying cell density to assess transformation efficiency. All libraries at this step yielded greater than 100,000 colonies corresponding to greater than 30× coverage for perfect mutations assuming 0.3% of the library has indels. All libraries (corresponding to different subpools) of a given target gene were pooled together at an equimolar ratio, resulting in a mixture of mutations for every amino acid position (Supplemental Figure S3C). This completes a single mutation library.

### Double mutations library generation

The double mutation library was generated by using the single mutation library as the target gene backbone for the insertion of another oligo subpool. Each oligo subpool was repeated using the methods described above (SPINE method) and mixed with equimolar ratio. This results in double mutations only across gene fragments and not within fragments. For the high number of variants expected, the Golden Gate reaction was transformed in E. cloni®10G ELITE electrocompetent cells (Lucigen). All libraries at this step yielded greater than 5,000,000 colonies corresponding to greater than 20× coverage.

### Bacterial Two-hybrid assay

The bacterial two-hybrid assay is based on PDZ3 binding to the CRIPT ligand. PDZ3 variants with a high affinity for the CRIPT ligand will recruit RNA polymerase α-subunit initiating expression of chloramphenicol acetyltransferase. This is a positive selection for highly functional PDZ3 variants. This system replicates the work of Salinas et al. ^37^ and all plasmid and cell reagents were received as a gift from Rama Ranganathan. The selection was performed with triplicate experiments. Plasmid from cells before selection and after selection was purified and the region covering the PDZ3 sequence was PCR amplified for 12 cycles with Illumina sequencing adapters. Amplicon DNA was purified with 1% agarose gel.

### NextGen Sequencing

Libraries were sequenced using Illumina MiSEQ in 150 bp paired-end configuration. Allele frequency for single mutation and double mutation was determined by joining paired sequences with bbmerge, trimming and filtering sequences with bbduk, and a custom python script to identify alleles only matching the OLS programmed mutation. Specifically, sequence alignment was performed by first joining paired sequences with bbmerge, trimming ends and filtering with bbduk and a custom python script to identify alleles only matching the programmed mutation in the OLS pools. The 150 bp paired-end sequences when joined together provide full coverage of the PDZ gene. This was done using bbmerge with the ‘xloose’ setting for strictness and a ‘minoverlap’ of 4 bp. This allows for greater number of reads to be merged for allele analysis. The 5’ extension setting at 2 bp allows for reads to be extended by 2 nucleotides for low overlap, but only allowing for 2 iterations (‘ecct extend2=2’). Merged reads were trimmed with bbduk with the literal string of the Illumina adpaters. The minimum adapter length was set to 7 bp to allow for incomplete Illumina adpaters (‘mink=7’) and quality trimming using Q10 and minimum length equal to PDS95PDZ3 gene length (249bp). Each processed read was then checked if it was the original sequence (recorded as WT), if not each read was analyzed for mutations at each position to search for mutations from the input library which were programmed on the OLS chip. If more mutations were found than expected (single or double) or if the read contained a mutation that did not match the programmed mutation it was removed and recorded as a bad read or a false positive, respectively. With read-pass filters that only recognize programmed mutations, we reduced the false-positive reads introduced by library generation and sequencing steps (Illumina reported at 1%). We detected and discarded on average 5% of reads due to false-positive mutations. Sequencing statistics are shown in **Supp. Fig. 2.1**. (single mutants) and **Supp. Fig. 3.1**. (double mutants).

## Data analysis

Read count data for all replicates (three biological replicates, 3 technical replicates) was summed (see supplemental information for all datasets).

### Fitness & Epistasis

Data analysis of read count data adapted workflow and scripts reported by Schmiedel at al. ^25^ with minor adaptations. Specifically, a 90% confidence interval (W_high_ and W_low_) was determined for single and double mutant fitness from read count data by using a Poisson distribution. Fitness confidence was calculated as

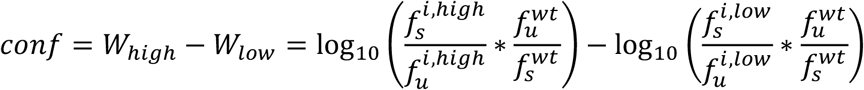

### flexddG Rosetta Backrub

Using PDB 1BE9 as the input structure, calculation of mutation effects on protein stability was implemented in RosettaScript as described by [ref] using Python scripts deposited at https://github.com/Kortemme-Lab/flex_ddG_tutorial. For each single and double mutant an ensemble of 35 mutant models were generated. Monte Carlo backrub was run for 35,000 steps. Rosetta energy scores are calculated using the Rosetta Talaris energy function refit with a generalized additive model ^34^.

## Acknowledgements

WCM is a Howard Hughes Medical Institute Gilliam Fellow and National Science Foundation Graduate Fellow. We thank Yungui He with providing support and reagents for the assays, and Mikael Elias for discussions. We also thank the U of MN Genomics Center for assistance with sequencing.

## Data Availability

Raw sequencing reads are deposited at https://www.ncbi.nlm.nih.gov/bioproject/642160

## Supplementary Materials for

## This PDF file includes

### Supplemental Figures

Supplemental Figure 1.1

Supplemental Figure 1.2

Supplemental Figure 2.1

Supplemental Figure 2.2

Supplemental Figure 2.3

Supplemental Figure 3.1

Supplemental Figure 3.2

Supplemental Figure 4.1

Supplemental Figure 4.2

Supplemental Figure 5.1

Supplemental Figure 6.1

**Supplemental Figure 1.1.**
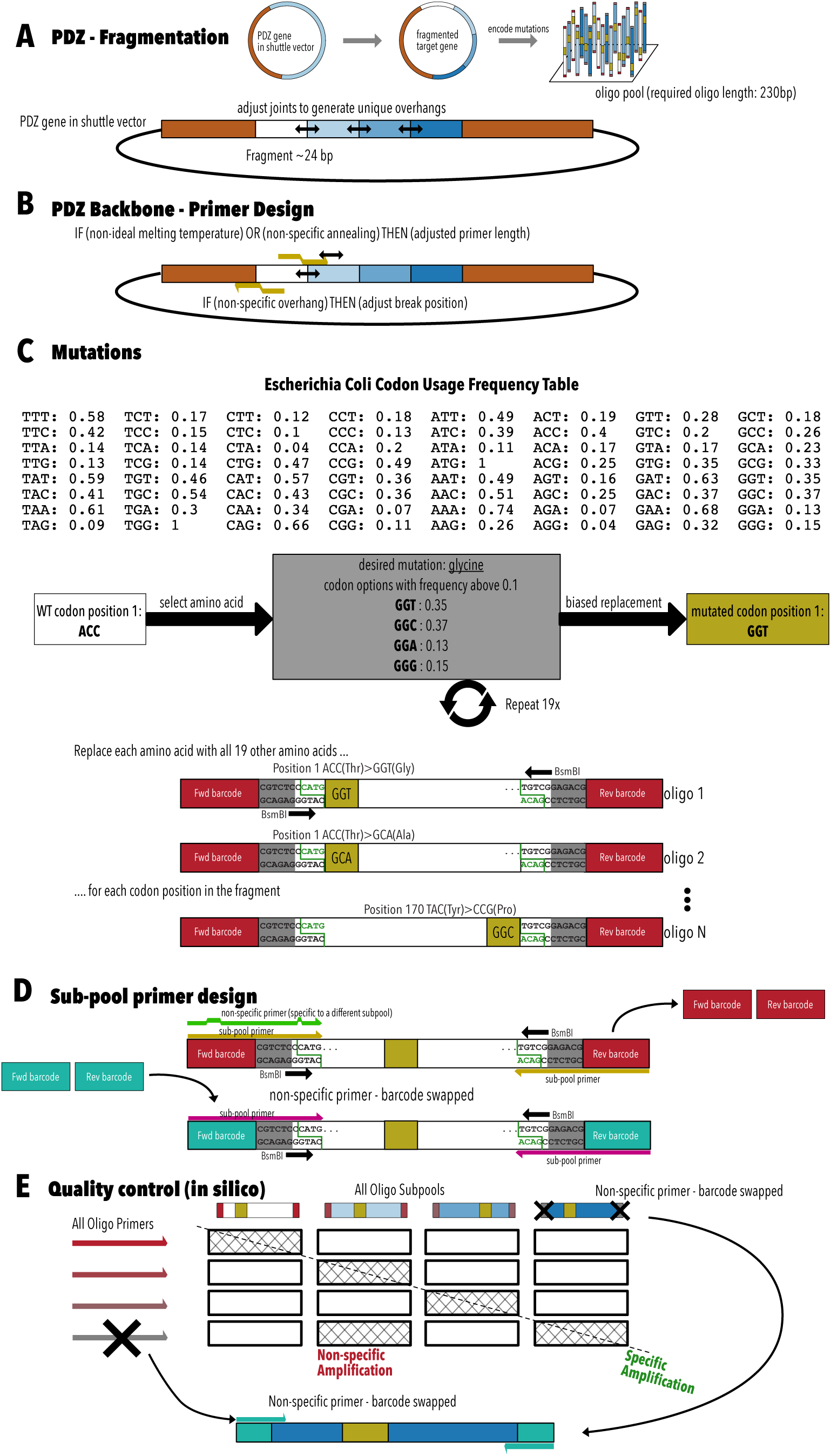
In silico design of oligos and primers. **A**, The PSD95 PDZ3 gene (within its shuttle vector) is fragmented into 10 fragments. Fragment break sites are adjusted for unique restriction enzyme cut overhangs. **B**,A set of gene primers are designed for each fragment for inverse PCR.These primers will amplify everything except the fragment and add an inward-facing BsmBI recognition site. **C**, An oligo pool is designed for each fragment and within the pool an oligo is designed for each amino acid within that fragment and for each of the 19 mutations. Each oligo consists of the fragment sequence it is replacing, sub-pool specific amplification barcodes, inward-facing BsmBI site that will match the cut site of the gene primers, and the mutation. **D**, To retrieve a specific sub-pool of oligos, primers are designed based on bio-orthogonal barcodes. This amplification is made specific by swapping barcodes until unique amplification is found. **E**, When combining the subpools from many genes, there is a chance of non-specific amplification. Quality control is performed on every oligo primer and oligo subpool for non-specific amplification. If found, the barcode is swapped for unique amplification.

**Supplemental Figure 1.2.**
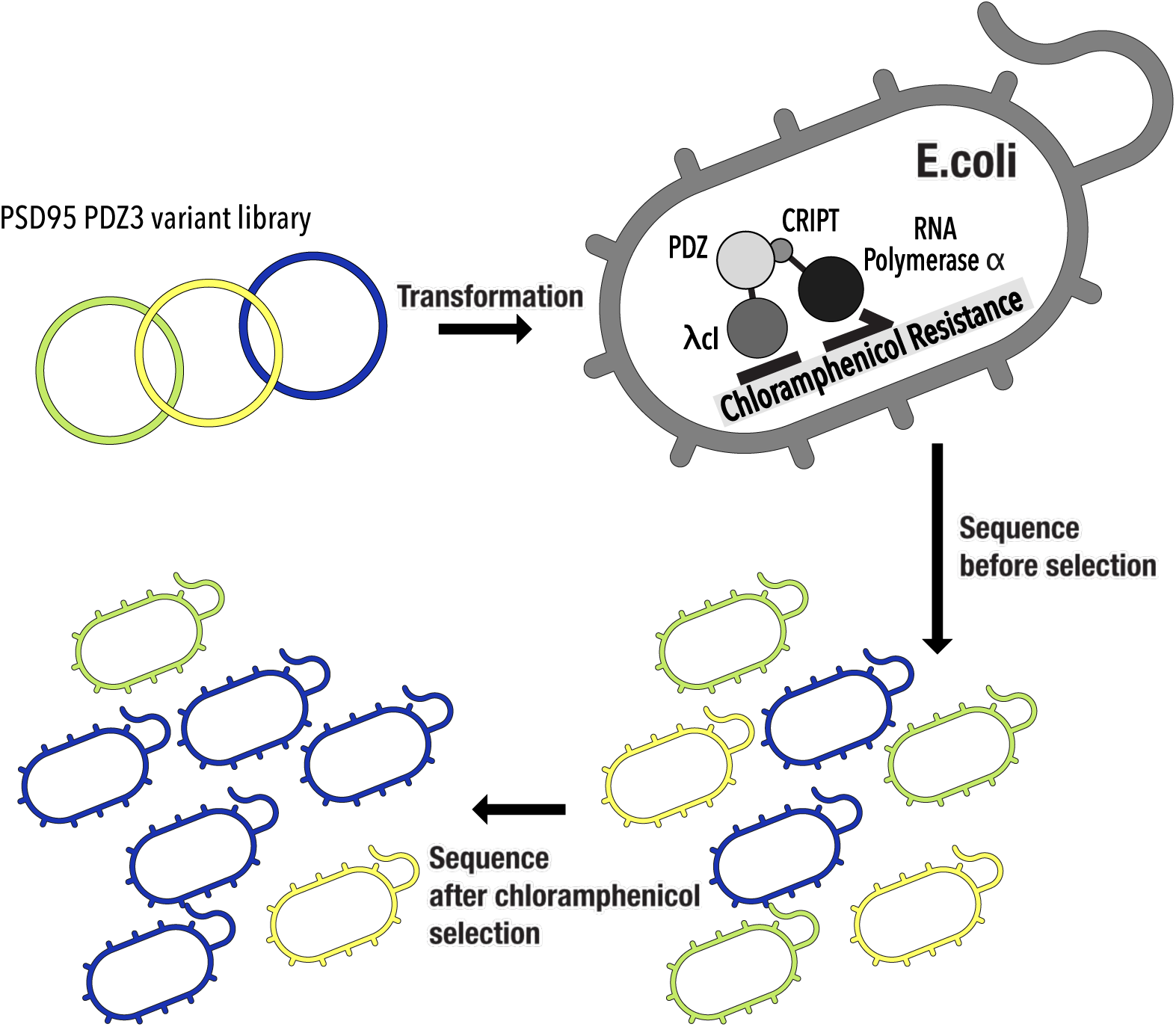
Bacterial two-hybrid itness assay. Mutant libraries are transformed into pZE1RM+pZA31+MC4100Z1 E. coli that have chromosomal copies of the lac repressor lacIQ and the tet repressor TetR. Each PDZ variant is fused to the λcl DNA binding domain and expressed under control of a lac promoter, while the CRIPT ligand is fused to the RNA polymerase α-subunit. When CRIPT ligand interacts with PDZ, chloramphenicol acetyltransferase is expressed, allowing the cell to survice challenge with the antibiotic chloramphenicol. By sequencing plasmid DNA isolated from transformed E.coli before and after chloramphenicol selection, the relative fitness of each variant can be calculated from read count data.

**Supplemental Figure 2.1.**
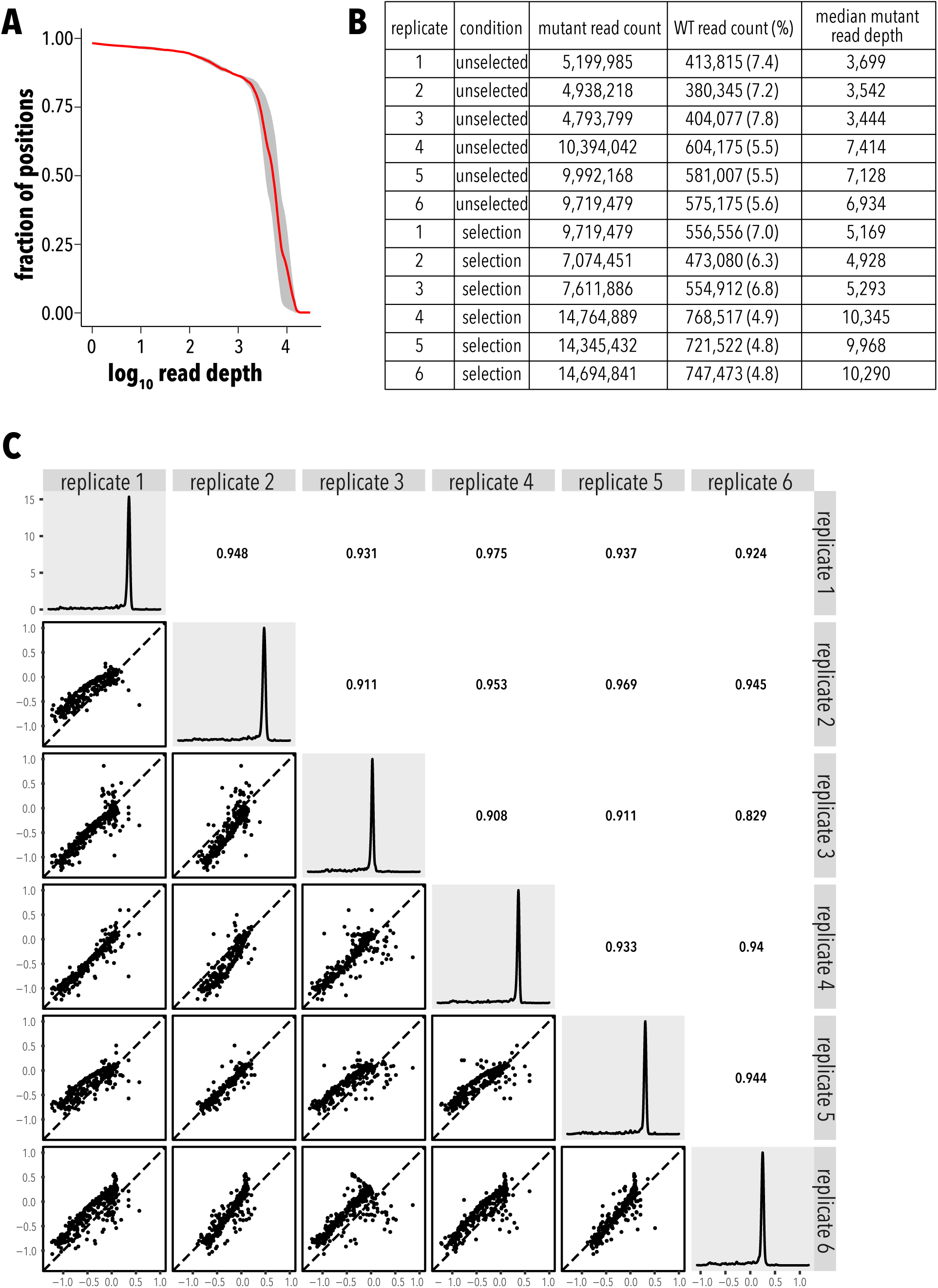
Single mutant dataset statistics. **A**, Mean read depth (red line, n=6) and confidence interval (shaded grey area). **B**, Count statistics. **C**, Fitness distribution for each replicate are shown on the diagonal. Replicate vs. replicates scatterplots are shown in the lower left triangle and Pearson correlation coefficients are shown in the upper right triangle.

**Supplemental Figure 2.2.**
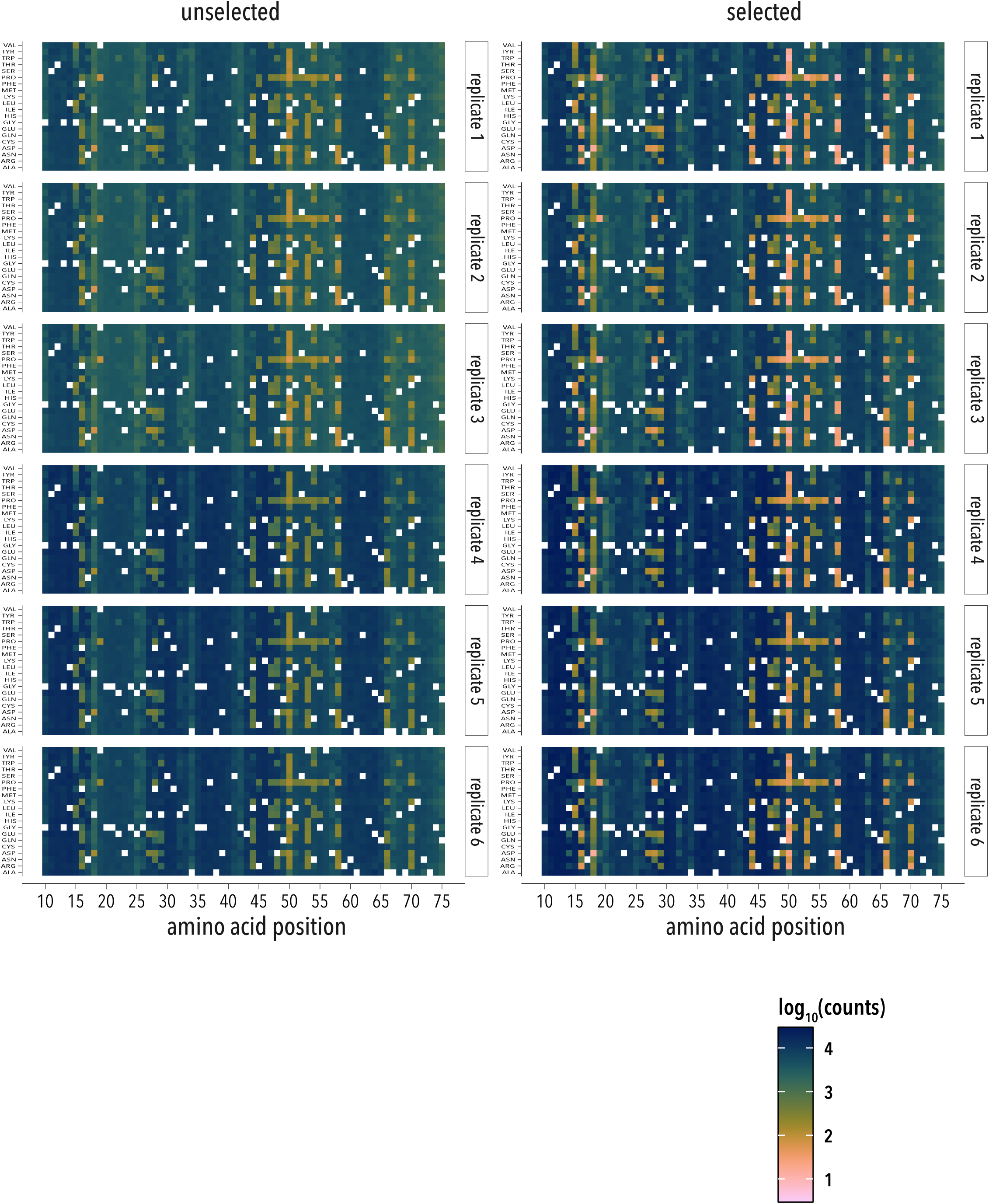
Single mutant library read count distribution by position and mutation for each replicate. Replicate where highly repeatable suggesting that underrepresented position and mutations are caused by sampling at the library construction stage.

**Supplemental Figure 2.3.**
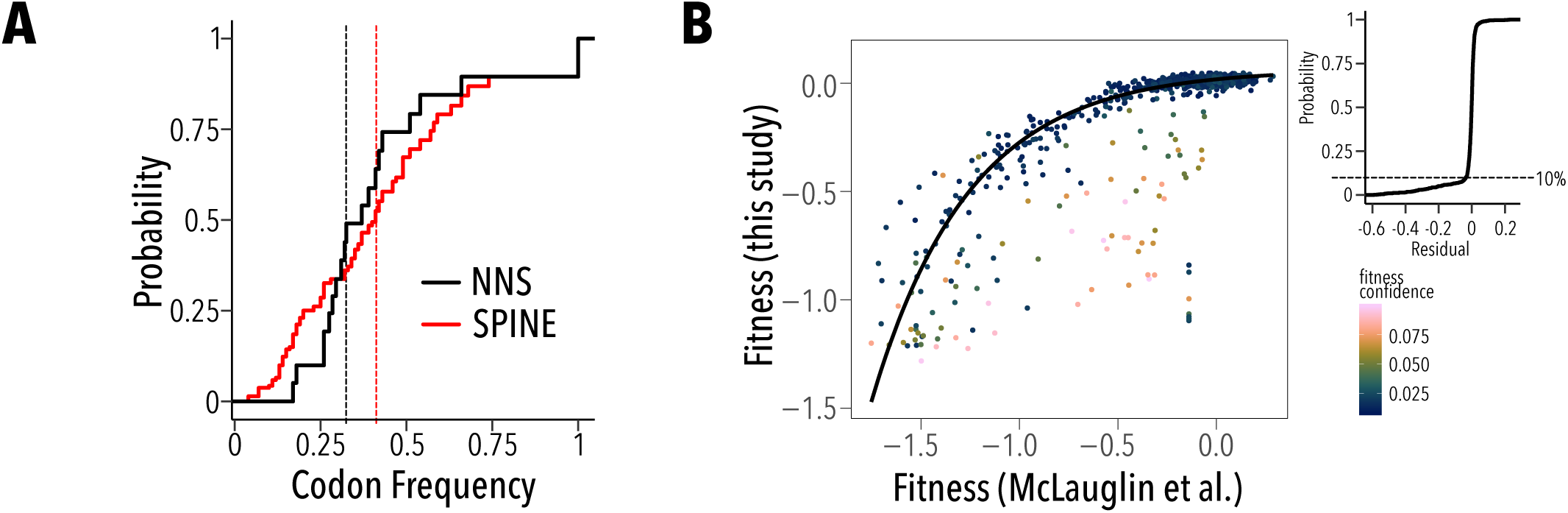
SPINE-generated libraries used optimal codons more often. **A**, Empirical cumulative distribution of codon frequency in a NNS degenerate codon library (black line) and the SPINE-generated library (red line) for PSD95 PDZ3. Vertical dashed line represent median usage frequency of optimal codons. With SPINE, more adapted codons are used more often. **B**, Comparing the fitness effect of single mutants in this study and McLaughlin et al. shows a monotonic, but non-linear relationship. Data points are colored by confidence in fitness determination, which is based on 90% Poisson confidence interval (see Methods). Data is fit to an exponential model (black line). he few (<10%) outlying residues (see inset empirical cumulative distribution function of fitting residuals) often had low fitness confidence.

**Supplemental Figure 3.1.**
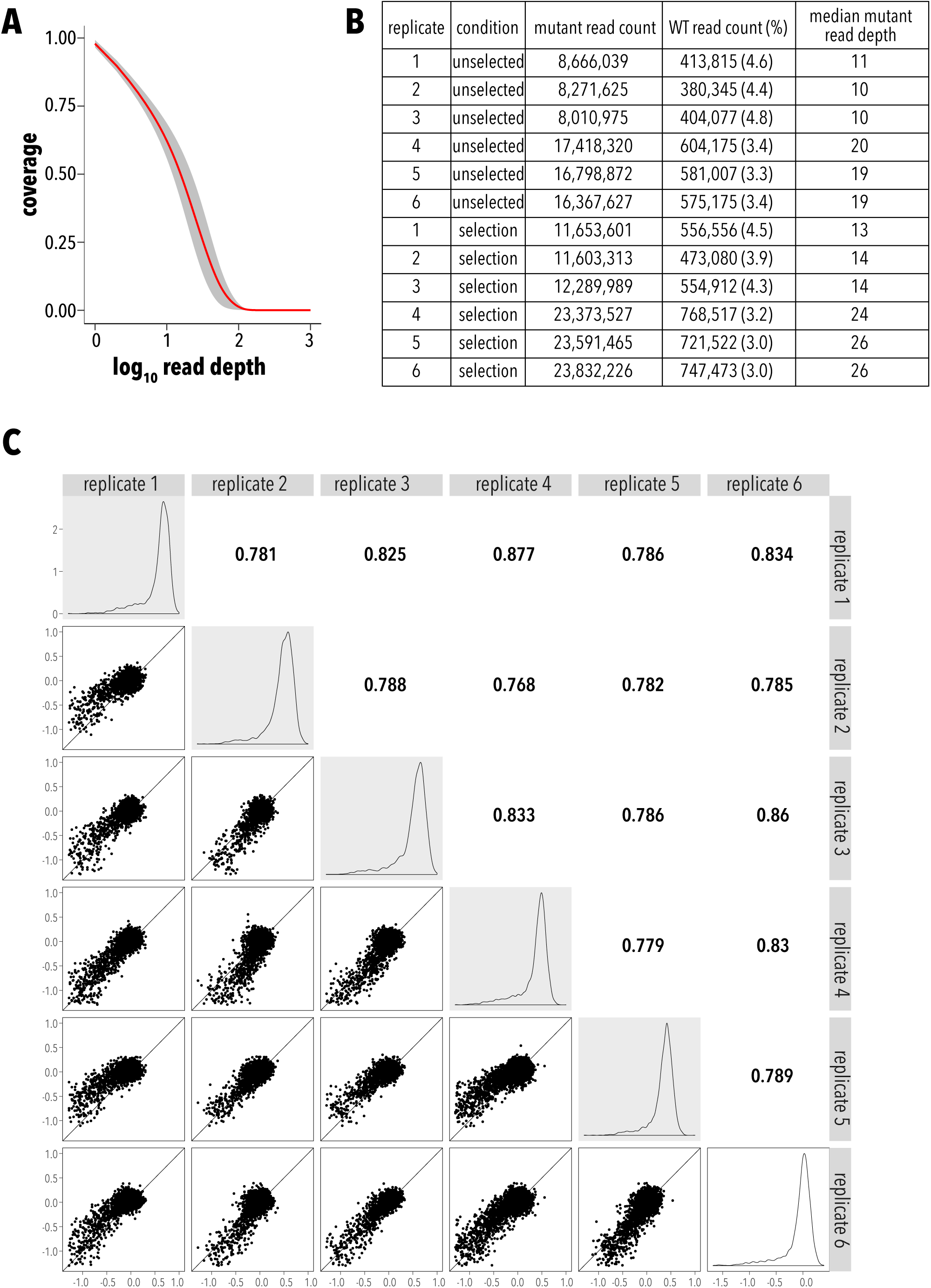
Double mutant dataset statistics. **A**, Mean read depth (red line, n=6) and confidence interval (shaded grey area). **B**, Count statistics. **C**, Fitness distribution for each replicate are shown on the diagonal. Replicate vs. replicates scatterplots are shown in the lower left triangle and Pearson correlation coefficients are shown in the upper right triangle.

**Supplemental Figure 3.2.**
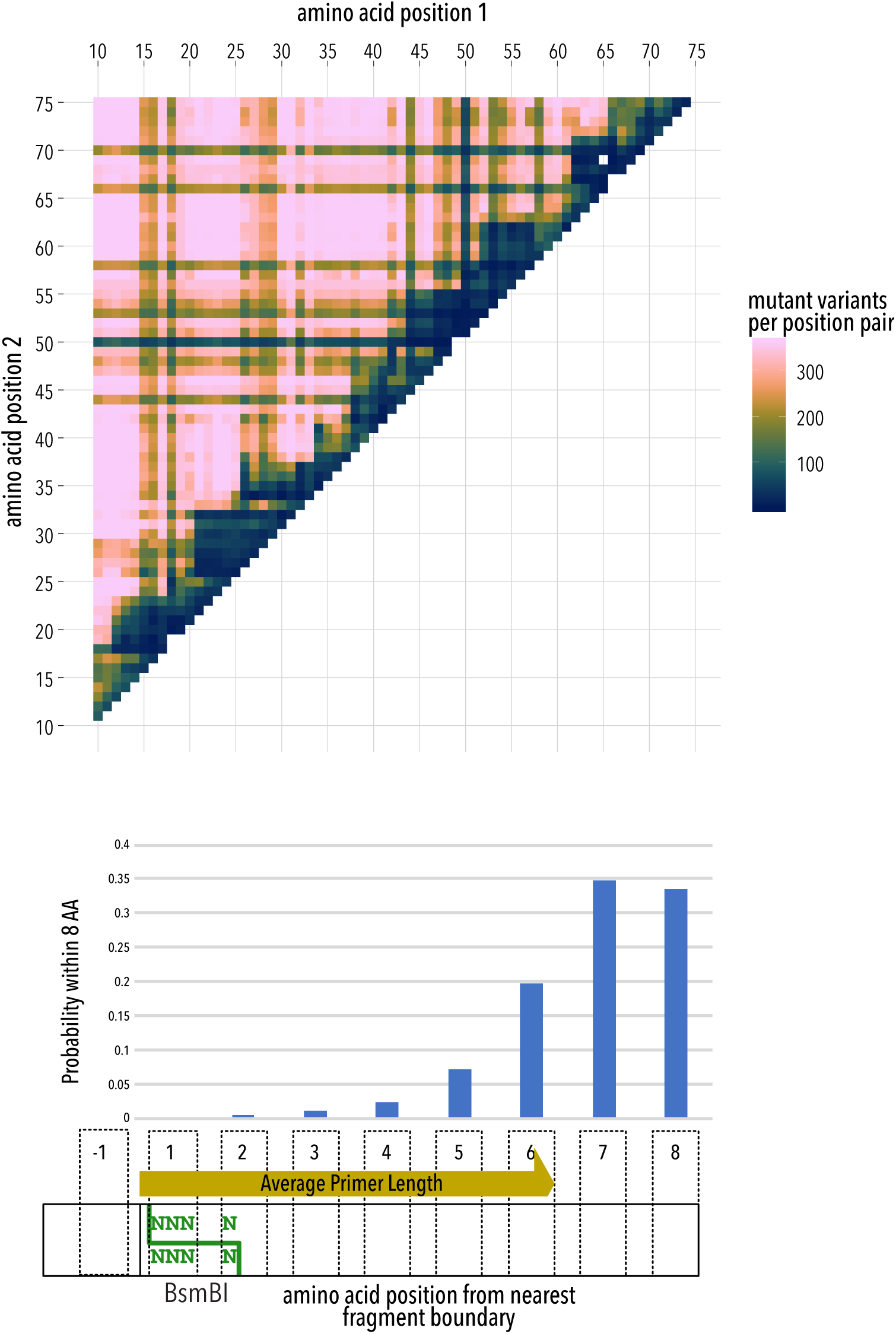
Double mutant library missing data. **A**, Positional map showing how many of the 19×19=361 possible mutations are represented in the count data after passing read quality threshold filters. Most position pairs with low coverage are <6 amino acids in linear sequence distance apart. **B**, This is due the nature of SPINE-mediate library assembly from oligo fragments. Nucleotides that are one position away are not possible due to the BsmBI cutsite, while pairs between connected fragments have an exponential increase in probability the further they are from the break site. This low probability for closely connected pairs is due to primer annealing and potential for mismatch bases.

**Supplemental Figure 4.1.**
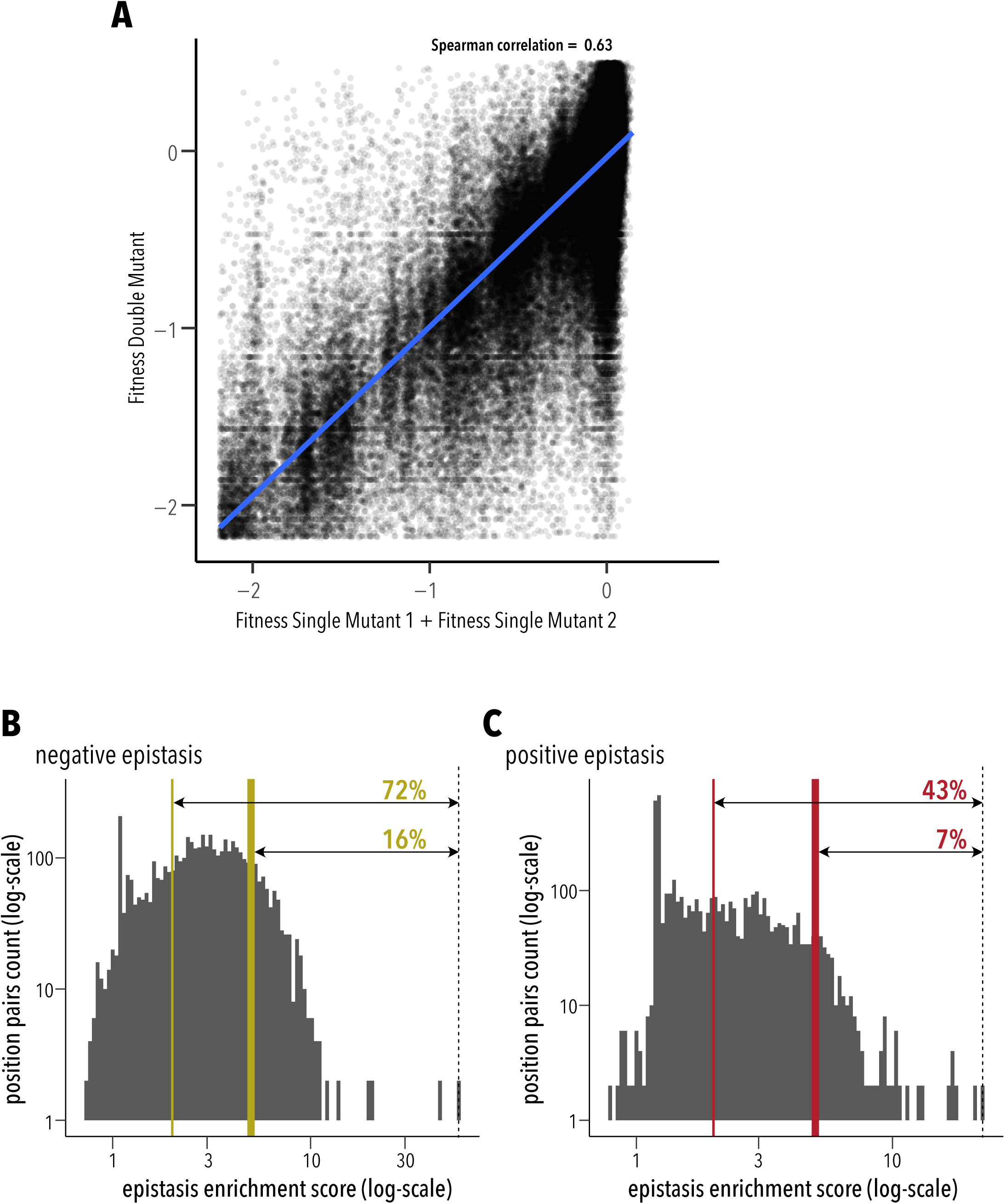
Prevalence of epistasis. **A**, The fitness of single mutants predicts double mutant fitness only moderately well (Spearman correlation coefficient 0.63) suggesting widespread deviation from expected additivity without epistasis. **B**, Negative epistasis with an enrichment score > 2 was observed in 72% of quantifiable position pairs (thin yellow line); stronger enrichment scores > 5 in 16% of quantifiable position pairs (thick yellow line). **C**, positive epistasis enrichment greater > 2 or > 5 was found in 43% (thin red line) or 7% (thick red line) of quantifiable position pairs, respectively.

**Supplemental Figure 4.2.**
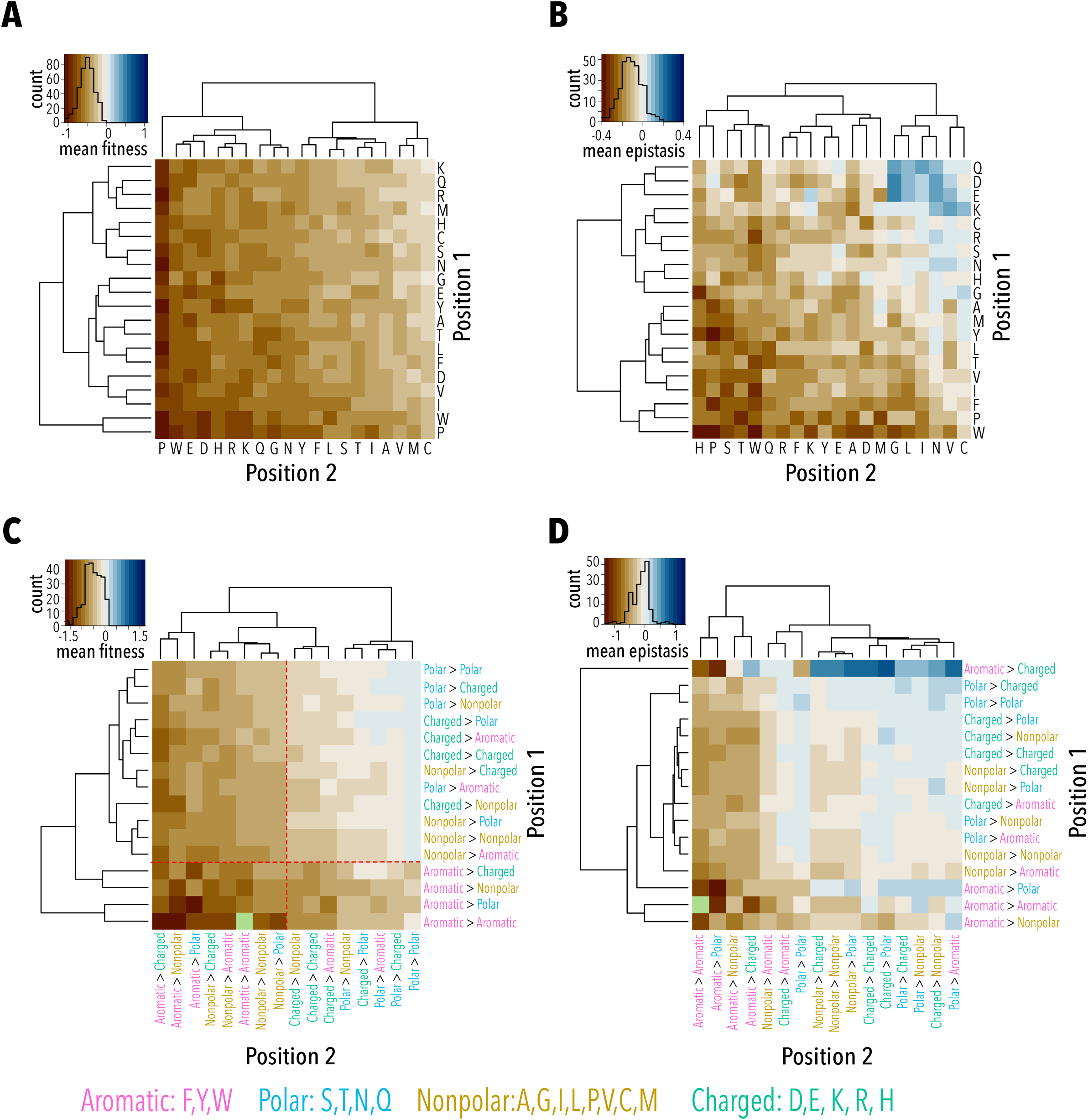
Physicochemical properties of mutation pairs and their role in itness and epistasis. **A**, Ordered heatmap of double mutant fitness grouped by mutant amino acid in either position. Fitness of double mutants is particularly impaired when both positions are mutated to disruptive (proline), bulky (tryptophan), or charged (glutamate, aspartate) amino acids. **B**, Ordered heatmap of epistasis in a double mutant grouped by mutant amino acid in either position. As expected, mutations to bulky aromatics or proline show strong negative epistasis in the background of proline and tryptophan mutations at a second site (exhausted excess stability). This is also true for polar and many charged residues. However, the same polar and charged residues in the background of small non-polar (valine, leucine, isoleucine) mutations show positive epistasis. **C**, Heatmap of double mutant fitness grouped by change in physicochemical properties. Fitness is strongly impaired when both wildtype position are aromatic or non-polar residues. **D**, Aromatic residues, in particular, show strong stratification with respect to physicochemical properties of the second site mutation hinting at the specifc nature of the underlying mechanisms.

**Supplemental Figure 5.1.**
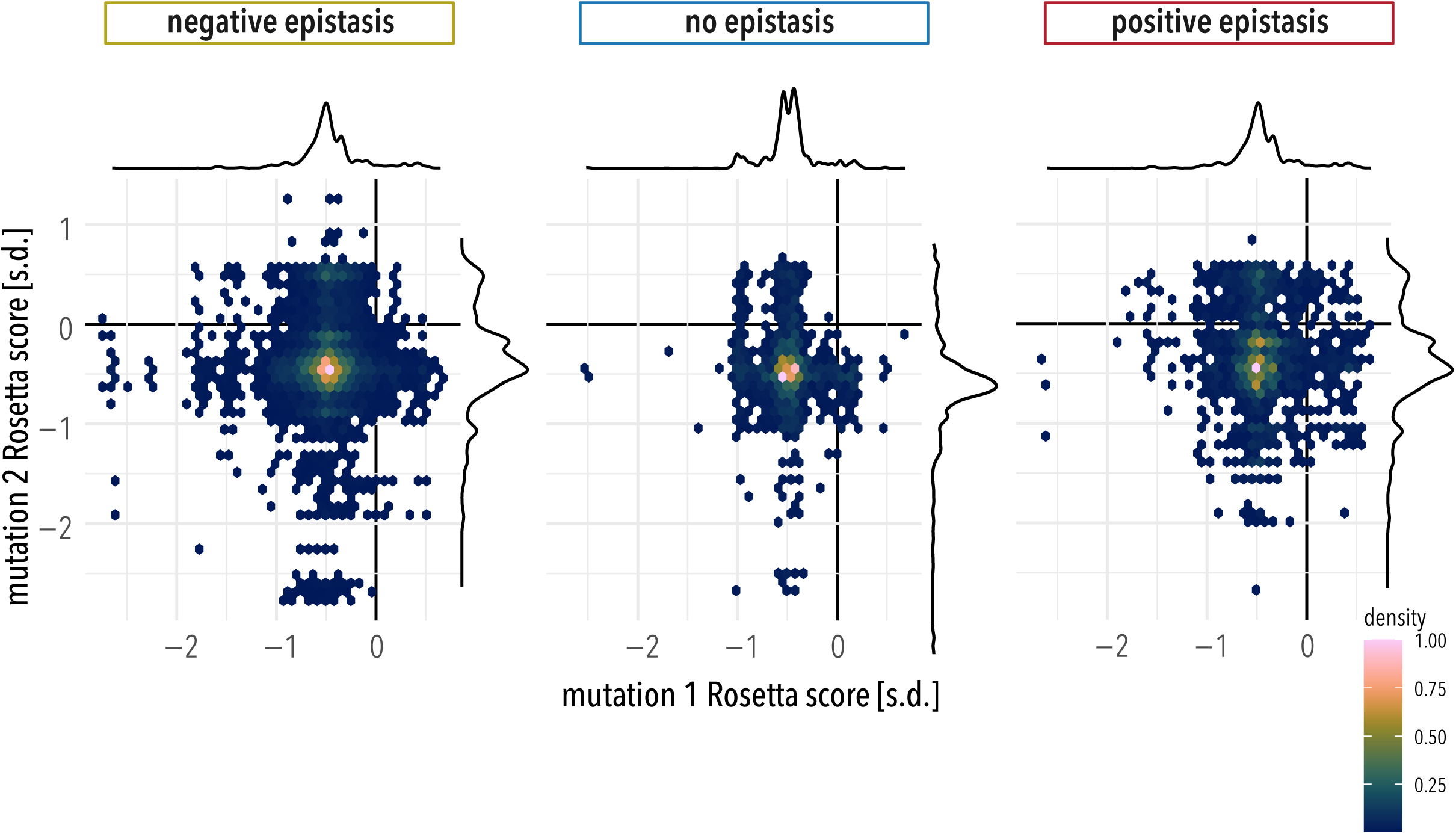
Distribution of z-scored Rosetta scores for single mutants in negative epistasis, no epistasis, and positive epistasis subsets. 2D histograms and marginal density plots of single mutant Rosetta scores (ΔΔG) for each double mutant.

**Supplemental Figure 6.1.**
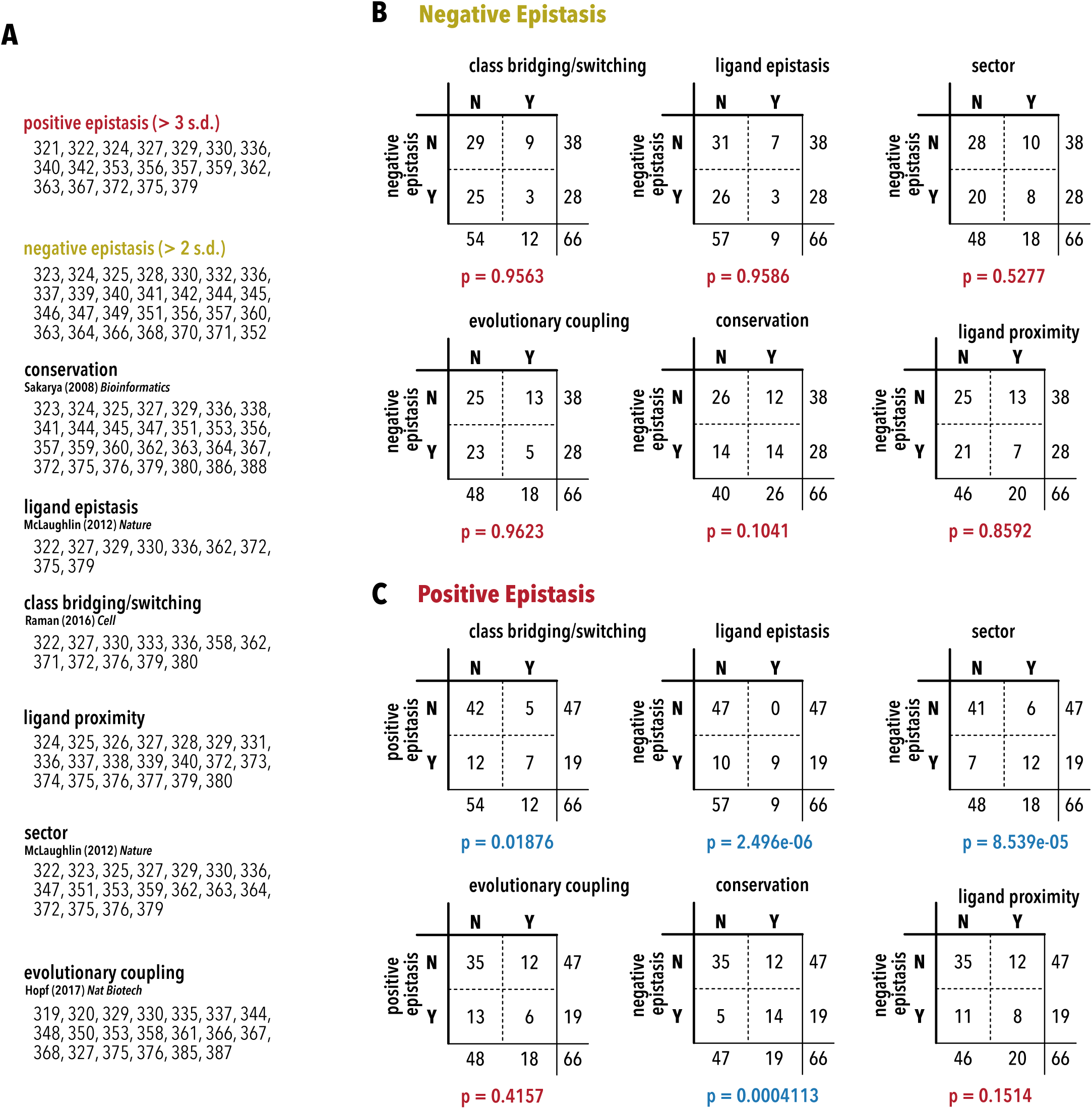
Contingency tables. **A**, Residue groupings. Independence of negative epistasis (**B**) and positive epistasis (**C**) in PSD95 PDZ3 with respect to different residue groupings was tested using Fisher’s Exact Test.

- The residue group that is class bridging / class switching (adaptive) was taken from Raman et al.
- The residue group that is sector positions, in proximity to ligand, or epistatic with respect to binding the wildtype CRIPT ligand vs. a class-switching T_-2_F mutant was taken from McLaughlin et al.
- Evolutionary coupling had been computed for DLG1 PDZ1 (residues 214-317, 40% identity, 68% similarity to PSD95 PDZ3) by Hopf et al. and deposited at https://marks.hms.harvard.edu/evmutation/index.html. Residues with evolutionary coupling score > 0.5 are shown. Residue numbering was establish from a structural alignment of hDLG-PDZ1 (PDB 3RL7) to PSD95 PDZ3 (PDB 1BE9).
- Conservation was calculated as using the Kullback-Leibler (KL) divergence of positional amino acid frequency in a PDZ family alignment reported by Sakaraya et al. versus the amino acid frequency in vertebrate proteins deposited in Uniprot. Residues with KL divergence > 1.5 are shown.

Negative epistasis was not correlated with any residue grouping; the null hypothesis of independence was not rejected (p > 0.5). Positive epistasis was enriched in class switching/bridging positions, ligand epistasis positions, sector positions, and conserved positions, but not position that are in proximity to the CRIPT ligand or those that are evolutionarily coupled.

## References

1. Fowler DM, Fields S. Deep mutational scanning: a new style of protein science. Nat Methods. 2014; 11(8) 801–807. doi: 10.1038/nmeth.3027. PubMed PMID: 25075907. PMCID: PMC4410700.

2. Firnberg E, Labonte JW, Gray JJ, Ostermeier M. A comprehensive, high-resolution map of a gene’s fitness landscape. Mol Biol Evol. 2014; 31(6) 1581–1592. doi: 10.1093/molbev/msu081. PubMed PMID: 24567513. PMCID: PMC4032126.

3. Fowler DM, Araya CL, Fleishman SJ et al. High-resolution mapping of protein sequence-function relationships. Nat Methods. 2010; 7(9) 741–746. doi: 10.1038/nmeth.1492. PubMed PMID: 20711194. PMCID: PMC2938879.

4. Melamed D, Young DL, Gamble CE, Miller CR, Fields S. Deep mutational scanning of an RRM domain of the Saccharomyces cerevisiae poly(A)-binding protein. RNA. 2013; 19(11) 1537–1551. doi: 10.1261/rna.040709.113. PubMed PMID: 24064791. PMCID: PMC3851721.

5. Mishra P, Flynn JM, Starr TN, Bolon DNA. Systematic Mutant Analyses Elucidate General and Client-Specific Aspects of Hsp90 Function. Cell Rep. 2016; 15(3) 588–598. doi: 10.1016/j.celrep.2016.03.046. PubMed PMID: 27068472. PMCID: PMC4838542.

6. Roscoe BP, Bolon DN. Systematic exploration of ubiquitin sequence, E1 activation efficiency, and experimental fitness in yeast. J Mol Biol. 2014; 426(15) 2854–2870. doi: 10.1016/j.jmb.2014.05.019. PubMed PMID: 24862281. PMCID: PMC4102620.

7. Starita LM, Pruneda JN, Lo RS et al. Activity-enhancing mutations in an E3 ubiquitin ligase identified by high-throughput mutagenesis. Proc Natl Acad Sci U S A. 2013; 110(14) E1263–72. doi: 10.1073/pnas.1303309110. PubMed PMID: 23509263. PMCID: PMC3619334.

8. Araya CL, Fowler DM, Chen W, Muniez I, Kelly JW, Fields S. A fundamental protein property, thermodynamic stability, revealed solely from large-scale measurements of protein function. Proc Natl Acad Sci USA. 2012; 109(42) 16858–16863. doi: 10.1073/pnas.1209751109. PubMed PMID: 23035249. PMCID: PMC3479514.

9. Sarkisyan KS, Bolotin DA, Meer MV et al. Local fitness landscape of the green fluorescent protein. Nature. 2016; 533(7603) 397–401. doi: 10.1038/nature17995. PubMed PMID: 27193686. PMCID: PMC4968632.

10. Melnikov A, Rogov P, Wang L, Gnirke A, Mikkelsen TS. Comprehensive mutational scanning of a kinase in vivo reveals substrate-dependent fitness landscapes. Nucleic Acids Res. 2014; 42(14) e112. doi: 10.1093/nar/gku511. PubMed PMID: 24914046. PMCID: PMC4132701.

11. Brenan L, Andreev A, Cohen O et al. Phenotypic Characterization of a Comprehensive Set of MAPK1/ERK2 Missense Mutants. Cell Rep. 2016; 17(4) 1171–1183. doi: 10.1016/j.celrep.2016.09.061. PubMed PMID: 27760319. PMCID: PMC5120861.

12. Matreyek KA, Starita LM, Stephany JJ et al. Multiplex assessment of protein variant abundance by massively parallel sequencing. Nat Genet. 2018; 50(6) 874–882. doi: 10.1038/s41588-018-0122-z. PubMed PMID: 29785012. PMCID: PMC5980760.

13. Starita LM, Young DL, Islam M et al. Massively Parallel Functional Analysis of BRCA1 RING Domain Variants. Genetics. 2015; 200(2) 413–422. doi: 10.1534/genetics.115.175802. PubMed PMID: 25823446. PMCID: PMC4492368.

14. Bershtein S, Segal M, Bekerman R, Tokuriki N, Tawfik DS. Robustness-epistasis link shapes the fitness landscape of a randomly drifting protein. Nature. 2006; 444(7121) 929–932. doi: 10.1038/nature05385. PubMed PMID: 17122770.

15. Breen MS, Kemena C, Vlasov PK, Notredame C, Kondrashov FA. Epistasis as the primary factor in molecular evolution. Nature. 2012; 490(7421) 535–538. doi: 10.1038/nature11510. PubMed PMID: 23064225.

16. Starr TN, Thornton JW. Epistasis in protein evolution. Protein Sci. 2016; 25(7) 1204–1218. doi: 10.1002/pro.2897. PubMed PMID: 26833806. PMCID: PMC4918427.

17. Olson CA, Wu NC, Sun R. A comprehensive biophysical description of pairwise epistasis throughout an entire protein domain. Curr Biol. 2014; 24(22) 2643–2651. doi: 10.1016/j.cub.2014.09.072. PubMed PMID: 25455030. PMCID: PMC4254498.

18. Halabi N, Rivoire O, Leibler S, Ranganathan R. Protein sectors: evolutionary units of three-dimensional structure. Cell. 2009; 138(4) 774–786. doi: 10.1016/j.cell.2009.07.038. PubMed PMID: 19703402. PMCID: PMC3210731.

19. Lockless SW, Ranganathan R. Evolutionarily conserved pathways of energetic connectivity in protein families. Science. 1999; 286(5438) 295–299. doi: 10.1126/science.286.5438.295. PubMed PMID: 10514373.

20. McLaughlin RN, Poelwijk FJ, Raman A, Gosal WS, Ranganathan R. The spatial architecture of protein function and adaptation. Nature. 2012; 491(7422) 138–142. doi: 10.1038/nature11500. PubMed PMID: 23041932. PMCID: PMC3991786.

21. Raman AS, White KI, Ranganathan R. Origins of Allostery and Evolvability in Proteins: A Case Study. Cell. 2016; 166(2) 468–480. doi: 10.1016/j.cell.2016.05.047. PubMed PMID: 27321669.

22. Rollins NJ, Brock KP, Poelwijk FJ et al. Inferring protein 3D structure from deep mutation scans. Nat Genet. 2019; 51(7) 1170–1176. doi: 10.1038/s41588-019-0432-9. PubMed PMID: 31209393.

23. Hopf TA, Ingraham JB, Poelwijk FJ et al. Mutation effects predicted from sequence co-variation. Nat Biotechnol. 2017; 35(2) 128–135. doi: 10.1038/nbt.3769. PubMed PMID: 28092658. PMCID: PMC5383098.

24. Toth-Petroczy A, Palmedo P, Ingraham J et al. Structured States of Disordered Proteins from Genomic Sequences. Cell. 2016; 167(1) 158-170.e12. doi: 10.1016/j.cell.2016.09.010. PubMed PMID: 27662088. PMCID: PMC5451116.

25. Schmiedel JM, Lehner B. Determining protein structures using deep mutagenesis. Nat Genet. 2019; 51(7) 1177–1186. doi: 10.1038/s41588-019-0431-x. PubMed PMID: 31209395.

26. Diss G, Lehner B. The genetic landscape of a physical interaction. Elife. 2018; 7 doi: 10.7554/eLife.32472. PubMed PMID: 29638215. PMCID: PMC5896888.

27. Hietpas R, Roscoe B, Jiang L, Bolon DN. Fitness analyses of all possible point mutations for regions of genes in yeast. Nat Protoc. 2012; 7 1382–1396. doi: 10.1038/nprot.2012.069. PubMed PMID: 22722372. PMCID: PMC3509169.

28. Kobori S, Yokobayashi Y. High-Throughput Mutational Analysis of a Twister Ribozyme. Angew Chem Int Ed Engl. 2016; 55(35) 10354–10357. doi: 10.1002/anie.201605470. PubMed PMID: 27461281. PMCID: PMC5113685.

29. Kitzman JO, Starita LM, Lo RS, Fields S, Shendure J. Massively parallel single-amino-acid mutagenesis. Nat Methods. 2015; 12(3) 203-6, 4 p following 206. doi: 10.1038/nmeth.3223. PubMed PMID: 25559584. PMCID: PMC4344410.

30. Kosuri S, Eroshenko N, LeProust EM et al. Scalable gene synthesis by selective amplification of DNA pools from high-fidelity microchips. Nature biotechnology. 2010; 28(12) 1295.

31. LeProust EM, Peck BJ, Spirin K et al. Synthesis of high-quality libraries of long (150mer) oligonucleotides by a novel depurination controlled process. Nucleic Acids Res. 2010; 38(8) 2522–2540. doi: 10.1093/nar/gkq163. PubMed PMID: 20308161. PMCID: PMC2860131.

32. Coyote-Maestas W, Nedrud D, Okorafor S, He Y, Schmidt D. Targeted insertional mutagenesis libraries for deep domain insertion profiling. Nucleic Acids Res. 2019; doi: 10.1093/nar/gkz1110. PubMed PMID: 31745561.

33. Sakarya O, Conaco C, Egecioglu O, Solla SA, Oakley TH, Kosik KS. Evolutionary expansion and specialization of the PDZ domains. Mol Biol Evol. 2010; 27(5) 1058–1069. doi: 10.1093/molbev/msp311. PubMed PMID: 20026484.

34. Barlow KA, Ó Conchúir S, Thompson S et al. Flex ddG: Rosetta Ensemble-Based Estimation of Changes in Protein-Protein Binding Affinity upon Mutation. J Phys Chem B. 2018; 122(21) 5389–5399. doi: 10.1021/acs.jpcb.7b11367. PubMed PMID: 29401388. PMCID: PMC5980710.

35. Ellington A, Pollard Jr JD. Introduction to the synthesis and purification of oligonucleotides. Current Protocols in Nucleic Acid Chemistry. 2000; 1) A. 3C. 1-A. 3C. 22.

36. Hecker KH, Rill RL. Error analysis of chemically synthesized polynucleotides. Biotechniques. 1998; 24(2) 256–260. doi: 10.2144/98242st01. PubMed PMID: 9494726.

37. Salinas VH, Ranganathan R. Coevolution-based inference of amino acid interactions underlying protein function. Elife. 2018; 7 doi: 10.7554/eLife.34300. PubMed PMID: 30024376. PMCID: PMC6117156.

38. Kudla G, Murray AW, Tollervey D, Plotkin JB. Coding-sequence determinants of gene expression in Escherichia coli. Science. 2009; 324(5924) 255–258. doi: 10.1126/science.1170160. PubMed PMID: 19359587. PMCID: PMC3902468.

39. Domingo J, Baeza-Centurion P, Lehner B. The Causes and Consequences of Genetic Interactions (Epistasis). Annu Rev Genomics Hum Genet. 2019; 20 433–460. doi: 10.1146/annurev-genom-083118-014857. PubMed PMID: 31082279.

40. Domingo J, Diss G, Lehner B. Pairwise and higher-order genetic interactions during the evolution of a tRNA. Nature. 2018; 558(7708) 117–121. doi: 10.1038/s41586-018-0170-7. PubMed PMID: 29849145. PMCID: PMC6193533.

41. Guy MP, Young DL, Payea MJ et al. Identification of the determinants of tRNA function and susceptibility to rapid tRNA decay by high-throughput in vivo analysis. Genes Dev. 2014; 28(15) 1721–1732. doi: 10.1101/gad.245936.114. PubMed PMID: 25085423. PMCID: PMC4117946.

42. Canals R, Chaudhuri RR, Steiner RE et al. The fitness landscape of the African Salmonella Typhimurium ST313 strain D23580 reveals unique properties of the pBT1 plasmid. PLoS Pathog. 2019; 15(9) e1007948. doi: 10.1371/journal.ppat.1007948. PubMed PMID: 31560731. PMCID: PMC6785131.

43. Puchta O, Cseke B, Czaja H, Tollervey D, Sanguinetti G, Kudla G. Network of epistatic interactions within a yeast snoRNA. Science. 2016; 352(6287) 840–844. doi: 10.1126/science.aaf0965. PubMed PMID: 27080103. PMCID: PMC5137784.

44. Eriksson AE, Baase WA, Zhang XJ et al. Response of a protein structure to cavity-creating mutations and its relation to the hydrophobic effect. Science. 1992; 255(5041) 178–183. doi: 10.1126/science.1553543. PubMed PMID: 1553543.

45. Kellis JT, Nyberg K, Fersht AR. Energetics of complementary side-chain packing in a protein hydrophobic core. Biochemistry. 1989; 28(11) 4914–4922. doi: 10.1021/bi00437a058. PubMed PMID: 2669964.

46. Richards FM, Lim WA. An analysis of packing in the protein folding problem. Q Rev Biophys. 1993; 26(4) 423–498. doi: 10.1017/s0033583500002845. PubMed PMID: 8058892.

47. Bloom JD, Labthavikul ST, Otey CR, Arnold FH. Protein stability promotes evolvability. Proc Natl Acad Sci U S A. 2006; 103(15) 5869–5874. doi: 10.1073/pnas.0510098103. PubMed PMID: 16581913. PMCID: PMC1458665.

48. Otwinowski J, McCandlish DM, Plotkin JB. Inferring the shape of global epistasis. Proc Natl Acad Sci U S A. 2018; 115(32) E7550–E7558. doi: 10.1073/pnas.1804015115. PubMed PMID: 30037990. PMCID: PMC6094095.

49. Stiffler MA, Hekstra DR, Ranganathan R. Evolvability as a function of purifying selection in TEM-1 β-lactamase. Cell. 2015; 160(5) 882–892. doi: 10.1016/j.cell.2015.01.035. PubMed PMID: 25723163.

50. McCandlish DM, Shah P, Plotkin JB. Epistasis and the Dynamics of Reversion in Molecular Evolution. Genetics. 2016; 203(3) 1335–1351. doi: 10.1534/genetics.116.188961. PubMed PMID: 27194749. PMCID: PMC4937490.

51. Morgan RD, Luyten YA. Rational engineering of type II restriction endonuclease DNA binding and cleavage specificity. Nucleic Acids Res. 2009; 37(15) 5222–5233. doi: 10.1093/nar/gkp535. PubMed PMID: 19567736. PMCID: PMC2731914.

52. Poelwijk FJ, de Vos MG, Tans SJ. Tradeoffs and optimality in the evolution of gene regulation. Cell. 2011; 146(3) 462–470. doi: 10.1016/j.cell.2011.06.035. PubMed PMID: 21802129.

53. Reynolds KA, McLaughlin RN, Ranganathan R. Hot spots for allosteric regulation on protein surfaces. Cell. 2011; 147(7) 1564–1575. doi: 10.1016/j.cell.2011.10.049. PubMed PMID: 22196731. PMCID: PMC3414429.

54. Pincus D, Pandey JP, Creixell P, Resnekov O, Reynolds KA. Evolution and engineering of allosteric regulation in protein kinases. BioRxiv. 2017; 189761.

55. Rosensweig C, Reynolds KA, Gao P et al. An evolutionary hotspot defines functional differences between CRYPTOCHROMES. Nat Commun. 2018; 9(1) 1138. doi: 10.1038/s41467-018-03503-6. PubMed PMID: 29556064. PMCID: PMC5859286.

56. Sugimoto N, Nakano S-i, Yoneyama M, Honda K-i. Improved thermodynamic parameters and helix initiation factor to predict stability of DNA duplexes. Nucleic acids research. 1996; 24(22) 4501–4505.

57. SantaLucia Jr J, Hicks D. The thermodynamics of DNA structural motifs. Annu Rev Biophys Biomol Struct. 2004; 33 415–440.

58. Xu Q, Schlabach MR, Hannon GJ, Elledge SJ. Design of 240,000 orthogonal 25mer DNA barcode probes. Proc Natl Acad Sci U S A. 2009; 106(7) 2289–2294. doi: 10.1073/pnas.0812506106. PubMed PMID: 19171886. PMCID: PMC2631075.

59. Engler C, Kandzia R, Marillonnet S. A one pot, one step, precision cloning method with high throughput capability. PLoS One. 2008; 3(11) e3647. doi: 10.1371/journal.pone.0003647. PubMed PMID: 18985154. PMCID: PMC2574415.

